# Integration host factor regulates colonization factors in the bee gut symbiont *Frischella perrara*

**DOI:** 10.1101/2021.12.06.471457

**Authors:** K. Schmidt, G. Santos-Matos, S. Leopold-Messer, Y. El-Chazli, O. Emery, T. Steiner, J. Piel, P. Engel

## Abstract

Bacteria colonize specific niches in the animal gut. However, the genetic basis of these associations is often unclear. The proteobacterium *Frischella perrara* is a widely distributed gut symbiont of honey bees. It colonizes a specific niche in the hindgut and causes a characteristic melanization response. Genetic determinants required for the establishment of this association, or its relevance for the host, are unknown. Here, we independently isolated three point mutations in genes encoding the DNA-binding protein integration host factor (IHF) in *F. perrara*. These mutants abolished the production of an aryl polyene metabolite causing the yellow colony morphotype of *F. perrara*. Inoculation of microbiota-free bees with one of the mutants drastically decreased gut colonization of *F. perrara*. Using RNAseq we found that IHF affects the expression of potential colonization factors, including genes for adhesion (Type 4 pili), interbacterial competition (Type 6 secretion systems), and secondary metabolite production (colibactin and aryl polyene biosynthesis). Gene deletions of these components revealed different colonization defects depending on the presence of other bee gut bacteria. Interestingly, one of the T6SS mutants did not induce the scab phenotype anymore, despite colonizing at high levels, suggesting an unexpected role in bacteria-host interaction. IHF is conserved across many bacteria and may also regulate host colonization in other animal symbionts.

## Introduction

The digestive tract of many animals is colonized by specialized gut symbionts that occupy distinct physical niches and utilize diverse nutrients. Owing to the availability of genetic tools, gnotobiotic animal models and multi-omics approaches, we can now study the genetic features that allow gut symbionts to colonize various animal hosts including mammals, fishes, and insects (1–10).

The Western honey bee, *Apis mellifera,* is a particularly interesting model to characterize colonization factors of bacterial gut symbionts, due to its agricultural importance and the tractability of its gut microbiota. Honey bees harbor relatively simple yet highly specialized gut microbiota composed of 8-10 bacterial genera (11, 12). The wide distribution of these communities across social bees suggests long evolutionary associations with the host (13). Moreover, different members of the bee microbiota colonize distinct physical niches along the gut. Lactobacilli and Bifidobacteria predominate in the posterior hindgut (rectum), while Gammaproteobacteria (*Frischella perrara* and *Gilliamella* species) and a Betaproteobacterium (*Snodgrassella alvi*) preferentially colonize the anterior hindgut (ileum and adjacent pylorus, i.e., the transition zone between midgut and ileum) (11, 14–16). The partitioning of these bacteria into distinct gut compartments suggests the existence of specific bacterial and/or host mechanisms that facilitate colonization.

Most bacteria of the bee gut microbiota can be cultured and experiments with gnotobiotic bees have been established (17–21). Moreover, genomic analyses have provided important insights about the functional potential of bee gut symbionts and their adaptation to the gut environment (21–29). Yet, little is known about which genes are directly involved in establishing colonization in the bee gut and how these genes are regulated. The only symbiont that has been extensively studied in this respect is *S. alvi.* Using transposon sequencing and transcriptome analysis, Powell et al. determined genome-wide host colonization factors in *S. alvi* (7). Most genes with strong fitness effects were found to belong to three major categories: extracellular interactions, metabolism, and stress response. In particular, genes for attachment and biofilm formation were highly beneficial for colonization, which is in agreement with the observation that this gut symbiont adheres to the host epithelium of the ileum and forms a multispecies biofilm with *Gilliamella*.

The honey bee gut symbiont *Frischella perrara* belongs to the recently described family Orbaceae within the Gammaproteobacteria (30). It is taxonomically close to the bee gut symbiont *Gilliamella* and has a similar metabolism (31). However, compared to the other members of the bee gut microbiota*, F. perrara* shows a rather distinctive colonization phenotype. While *S. alvi* and *Gilliamella* have been reported to colonize the host epithelium of the entire ileum (32, 33), *F. perrara* preferentially colonizes the transition zone between the midgut and the ileum, i.e., the pylorus. Moreover, colonization with *F. perrara* leads to the appearance of a brown to black material on the luminal side of the epithelial surface between the cuticle layer of the host tissue and the adherent *F. perrara* cells (34). This so–called scab phenotype forms after 5-7 days post-colonization and has so far not been reported to be triggered by any other gut symbiont than *F. perrara*. Transcriptome analysis of the host showed that *F. perrara* elicits a specific immune response which includes the upregulation of the host melanization pathway likely responsible for the formation of the scab phenotype (17). *F. perrara* is highly prevalent across worker bees and colonies of *A. mellifera* (34, 35), and related bacteria have also been found in *Apis cerana* (36). Moreover, between 25%-80% of all worker bees of a colony harbor a visible scab phenotype in the pylorus region of the gut, which has been shown to strongly correlate with a high abundance of *F. perrara* (34). However, the impact of these phenotype on the host has remained elusive.

Genome sequencing of the type strain of *F. perrara* and comparison with other genomes of the Orbaceae family revealed the presence of several genomic islands that may be involved in the specific interaction of *F. perrara* with the host (37). These include a biosynthetic gene cluster for the production of the genotoxic metabolite colibactin (Clb), two distinct Type VI secretion systems (T6SSs) and associated effector proteins, Type I secretion systems, and fimbrial low-molecular-weight protein (Flp) pili genes. However, currently no genetic tools are available for *F*. *perrara* precluding studies about the role of these genetic factors in gut colonization or the induction of the scab phenotype.

Here, we report the isolation of a spontaneous mutant of *F. perrara* that possesses a strong colonization defect *in vivo*. Resequencing of the mutant revealed a single nonsynonymous point mutation in the gene encoding the alpha subunit of the DNA-binding protein integration host factor (IHF). Using a combination of gnotobiotic bee experiments, transcriptomics, and metabolite analyses, we characterized the genes regulated by IHF. We then established a gene deletion strategy for *F. perrara,* which allowed us to knockout some of the IHF-regulated genes and show that they impact gut colonization and scab development to different extent in the presence and absence of a complex community.

## Results

### Isolation of spontaneous IHF mutants affecting growth and colony morphology of *F. perrara*

Culturing *F. perrara* type strain PEB0191 (38) on modified tryptone yeast glucose (mTYG) agar resulted in the formation of yellow colonies. However, we occasionally observed the appearance of larger white colonies among the yellow ones (**Figure 1A**). Restreaking white colonies on fresh mTYG agar usually resulted in yellow colonies again. However, three white colonies that we identified in independent experiments did not change their appearance anymore, suggesting that we had isolated stable ‘white’ variants of *F. perrara* PEB0191 (**Figure 1B, Figure 1 - figure supplement 1**). Genome sequencing of the white variants revealed the presence of three different non-synonymous point mutations in the genes encoding the integration host factor (IHF). IHF is a widely distributed DNA-binding protein consisting of the IhfA/B heterocomplex (39–41). Strikingly, two point mutations were identical to each other, but occurred in the different subunits of IHF (*ihfA* and *ihfB*) resulting in a proline to lysine change at amino acid position 82 and 83, respectively (**Figure 1C**). The third point mutation resulted in a lysine to serine change at position 38 of IhfA (**Figure 1C**). Homology modelling showed that these amino acids are located in the region interacting with DNA, suggesting that the three mutations impact the DNA-binding properties of IHF (**Figure 1D**, (42)). As two of the isolated mutants occurred when generating gene deletions of *F. perrara,* they harbored additional genetic modifications (see Methods). Therefore, we focused further characterization of IHF on the mutation Pro83Lys (hereafter *ihfA\**) that occurred in the wild type (wt) background of *F. perrara* PEB0191. While the *ihfA\** strain consistently formed larger colonies than the wt strain on mTYG agar (**Figure 1A and 1B)**, there was no significant difference in growth in liquid culture (Permutation test, p = 0.097; **Figure 1 – figure supplement 2A**). However, light microscopy showed that cells of the mutant strain were on average slightly longer than cells of the wt (Kolgomorov-Smirnov test p<0.0001, **Figure 1 – figure Supplement 2B and C**).

**Figure 1.**
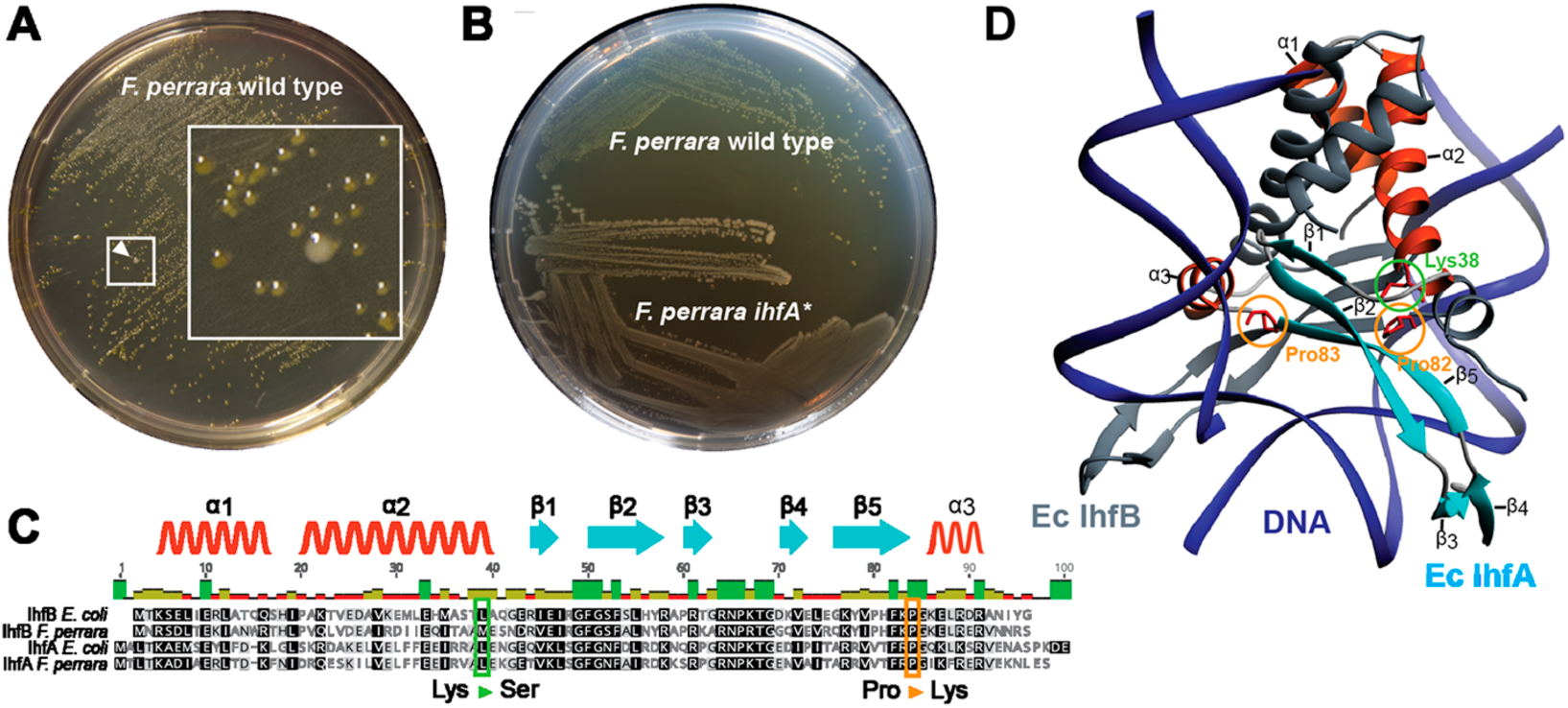
Isolation of a spontaneous *ihfA\** mutant of *F. perrara* displaying an altered colony morphotype. **(A)** Colonies of *F. perrara* PEB0191 (wt) after 48 hours of growth on mTYG agar. Arrowhead points at a larger white colony in between many yellow colonies. The area in the white square is magnified. (**B**) Colony morphology of *F. perrara* wt and the isolated white *ihfA\** mutant after growth on mTYG for 48 hours. (**C**) Protein sequence comparison of IhfA and IhfB of *F. perrara* wt and *E. coli* wt. The outlined positions refer to the residues mutated in the three spontaneous *ihfA* mutants: (i) lysine (Lys) to serine (Ser) at position 38 of *F. perrara* IhfA, (ii) proline (Pro) to lysine (Lys) at position 83 of *F. perrara* IhfA, (iii) proline (Pro) to lysine (Lys) at position 82 of *F. perrara* IhfB. Note that the numbers given on top of the alignment refer to alignment positions and not to positions in the individual sequences. Secondary structures are depicted above as ribbons (α-helix) and arrows (β-sheet) and are numbered according to their appearance in the protein and the structure shown in D. (**D)** Three-dimensional structure of *E. coli* IhfA/B heterocomplex with DNA (source protein databank NDB: PDT040). DNA is depicted in blue and IhfB in dark grey. IhfA is colored according to secondary structure: α-helix orange, β-sheet light blue and the rest in light grey. α-helices and β-sheets are numbered. The mutated Pro83 and Lys38 residues of *F. perrara* IhfA and the Pro82 residue of IhfB are marked with an orange and green circle, respectively.

### *F. perrara* produces an aryl polyene secondary metabolite that is responsible for the yellow colony morphotype

*F. perrara* PEB0191 encodes a genomic island that is homologous to aryl polyene (APE) biosynthetic gene clusters present in other Gammaproteobacteria (**Figure 2A**) (43). APEs are polyunsaturated carboxylic acids conferring a yellow pigmentation to bacterial cells (44, 45). To assess if *F. perrara* wt, but not the *ihfA\** mutant, produces an APE, we analyzed cell extracts of both strains by liquid chromatography coupled to heated electrospray ionization high-resolution mass spectrometry (HPLC-HES-HRMS). The data revealed a strongly UV-Vis-absorbent ion peak at *m/z* 323.1647 [M+H]^+^, which had a suggested molecular formula of C_21_H_23_O_3_ (**Figure 2B** and **2C**). In the *ihfA\** mutant, this ion was only present at trace amounts (**Figure 2C**). To characterize the metabolite in greater detail a larger pellet of *F. perrara* wt cultures was extracted and purified by several HPLC runs. Mass spectrometry- (MS) and UV-Vis-guided fractionation yielded an enriched extract that was analyzed by nuclear magnetic resonance (NMR) spectroscopy. The characteristic ions detected in MS-MS fragmentation experiments (**Figure 2 – figure supplement 1**), the UV-Vis spectrum with an absorption maximum at 415 nm (**Figure 2 – figure supplement 2**) in conjunction with NMR data (**Figure 2 – figure supplement 3-9, Supplementary Table 1**) suggest an aryl polyene structure identical to that reported in (43, 46) (**Figure 2D**). Unfortunately, it was not possible to connect the NMR substructures, because the central methines could not be assigned to chemical shifts (**Figure 2D**, **Supplementary Table 1)**. Comparison of the organic extracts of *F. perrara* wt and *E. coli* CFT073 provided further evidence that both produce the same compound (**Figure 2 – figure supplement 10**). Combined, these results suggest that the APE pathway is responsible for the yellow color of the wt colonies of *F. perrara* and is suppressed in the *ihfA\** mutant.

**Figure 2.**
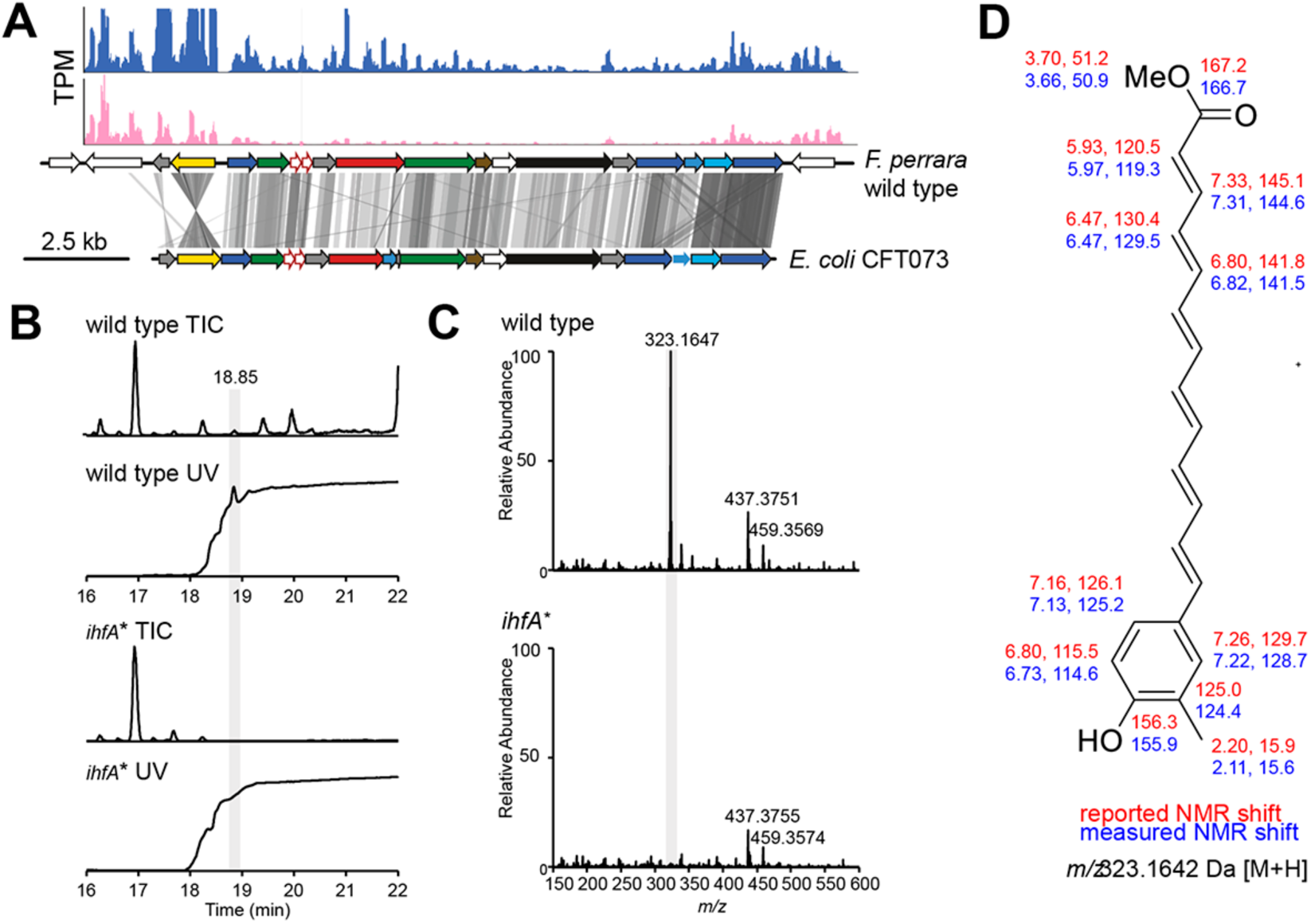
Metabolite analysis of *F. perrara* wt and the *ihfA\** mutant. **(A)** Comparison of gene synteny and sequence similarity of the genomic islands of *F. perrara* PEB0191 (top) and *E. coli* CFT073 (bottom) encoding the aryl polyene (APE) biosynthesis genes. Grey lines indicate homologous regions based on tblastx analysis. Plots were generated with genoplotR (82). Transcripts per million (TPMs) are shown on top of the genomic island for one RNAseq replicate of each *F. perrara* wt (blue) and the *ihfA\** mutant grown in vitro. Coverage plots were generated with the Integrated Genome Browser v9 (74). (**B)** Total ion chromatogram (TIC) and UV trace (λ=420 nm) of wt and *ihfA*\*. A peak highly abundant in the wt was discovered at 18.85 min. Its high UV absorbance at λ=420 nm indicated a conjugated carbon double bond system. (**C)** The normalized mass spectrum at 18.85 min reveals the ion *m/z* = 323.1647 Da to be approximately 50-fold more abundant in the wt compared to *ihfA*\*. (**D)** Enrichment of the ion containing fraction by HPLC followed by NMR experiments suggest a structure identical to that reported by (43). Reported (red) and observed (blue) ^1^H and ^13^C chemical shifts are shown. Central methines could not be assigned.

### The *ihfA\** mutant of *F. perrara* has a colonization defect and does not cause the scab phenotype

As APEs have been shown to increase protection from oxidative stress and contribute to biofilm formation (43, 46), we sought to test if the *ihfA\** mutation impacts bee gut colonization. We mono-associated microbiota-free bees with either *F. perrara* wt or *ihfA\**. Colonization with the wt strain resulted in a visible scab in 50% and 80% of all bees after five and ten days of colonization, respectively (n=18 and n=36 for both treatments for day 5 and day 10, respectively, **Figure 3A and 3B**). In contrast, none of the bees colonized with the *ihfA\** mutant developed a visible scab phenotype. To determine whether this difference was due to a general colonization defect of *ihfA*\*, we quantified the colonization levels of *F. perrara* at day 5 and day 10 post colonization using colony forming units (CFUs). While there was a trend towards lower colonization levels (fewer CFUs and more bees without detectable colonization) for the *ihfA\** mutant at day 5 post colonization, the difference was not statistically significant (**Figure 3C**, Wilcoxon rank-sum test p-value=0.076). However, at day 10 post-colonization, bees colonized with the wt strain showed significantly higher CFUs than the *ihfA\** mutant (**Figure 3C**, Wilcoxon rank-sum test p-value<0.0001). In fact, in 50% of all bees (n = 36) the colonization levels of *ihfA\** were below the detection limit of 500 CFUs (colony forming units) (**Figure 3C**).

**Figure 3.**
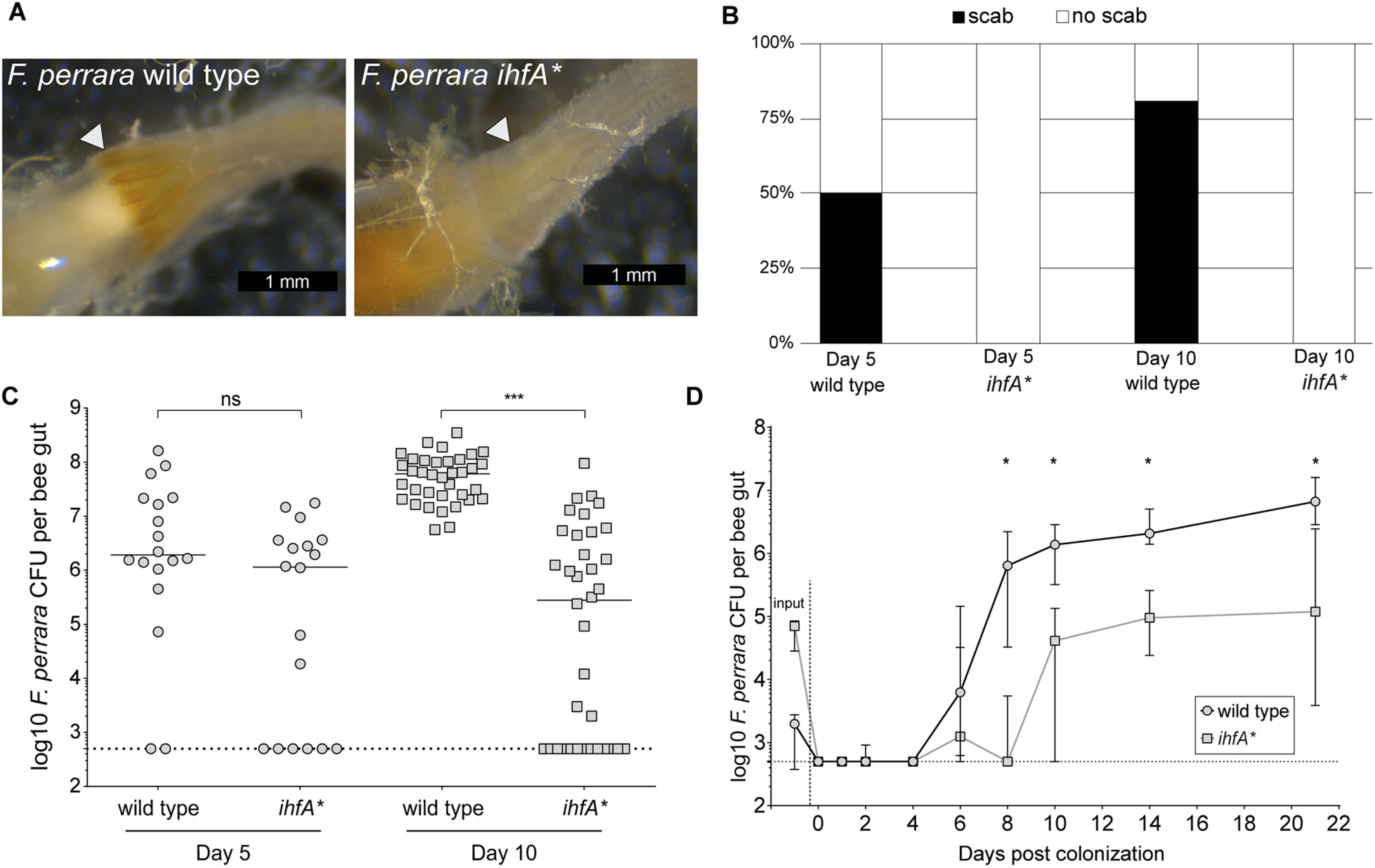
*F. perrara ihfA\** mutant displays a colonization defect. **(A)** Light microscopy pictures of pylorus region of bees colonized with *F. perrara* PEB0191wt or *ihfA\** 10 days post colonization. (B) Quantification of scab phenotype of bees 5 and10 days post colonization with n=18 and n=36 per treatment, respectively. (**C)** Quantification of colonization levels are measured by colony forming units (CFUs) at day 5 (n=18) and day 10 (n=36) post colonization. Wilcoxon rank-sum test was used to assess significant differences. (**D)** Time course experiment of bees colonized with *F. perrara* wt or *ihfA\**. Colonization levels were measured by CFUs every second day until day 10 and then at day 14 and day 21. n=12 bees per time point per treatment. Wilcoxon rank-sum test was used to assess significant differences per time point. Error bars represent median and interquartile range. Data from three independent experiments. *p<0.05, **p<0.01, ***p<0.001.

As the quantification of *F. perrara* was based on CFUs obtained for whole-gut tissue, we carried out a second experiment, in which we specifically assessed the colonization levels in the pylorus and the ileum region of the honey bee gut, using both CFUs and quantitative PCR (**Figure 3 - figure supplement 1**). The results were comparable to those obtained for the whole gut: at day 10 post colonization, there was a significant difference in the colonization levels of the wt and *ihfA\** in both the pylorus and the ileum (Wilcoxon rank-sum test p-value<0.05).

To obtain a better understanding of the colonization dynamics of *F. perrara* wt and the *ihfA\** mutant, we conducted a third gnotobiotic bee experiment in which we inoculated microbiota-free bees with one of the two strains and followed the colonization levels over 12 timepoints from day 0 (i.e. 4 h post inoculation) until day 22 post inoculation (**Figure 3D**). From the first time point at 4 h post inoculation until day 4 post inoculation, the bacterial levels were below the detection limit (i.e., below 500 CFUs) in both conditions. Between day 4 and day 8 post inoculation, the abundance of the wt increased rapidly to about 10^6^ CFUs per gut and then steadily further to 10^7^ CFUs per gut until the last time point. In contrast, the levels of the *ihfA\** mutant remained low until day 10 post colonization and reached on average no more than 10^5^ CFUs per gut until the last time point at day 22 post colonization. Notably, while we had used the same optical density of the two strains for colonizing microbiota-free bees, dilution plating revealed that there were fewer CFUs in the inocula for the wt compared to the *ihf\** mutant. Despite these differences the wt colonized much better than *ihfA\**. In summary, these results show that *ihfA\** has a strong colonization defect. It has a delayed colonization dynamics as compared to the wt, does not reach the same bacterial loads, and does not cause the scab phenotype, even though the bees were inoculated with more viable cells of *ihfA\** than the wt.

### Genes involved in symbiotic interactions are upregulated in *F. perrara* wt relative to the *ihfA\** mutant

IHF may not have a direct effect on gut colonization, but rather regulate the gene expression of host colonization factors. To test this, we assessed the transcriptional differences by RNA sequencing (RNA-seq) between the wt and *ihfA\** mutant when grown *in vitro*. We found that 358 out of 2,337 genes encoded in the genome of *F. perrara* were differentially expressed with a log2-fold change >|2| between the two strains (Fisher’s exact test with p<0.05 and FDR<5%). Of those, 237 and 121 genes were up- and down-regulated in *F. perrara* wt versus *ihfA*\*, respectively (**Figure 4A and 4B**, **Supplementary Dataset 1**). Among the genes upregulated in the wt, ‘Intracellular trafficking, secretion, and vesicular transport’ (COG U), ‘Extracellular structure’ (COG W), ‘Lipid transport and metabolism’ (COG I), ‘Mobilome: prophages and transposases’ (COG X), and ‘Secondary metabolites biosynthesis, transport and catabolism’ (COG Q) were significantly enriched (Fisher’s exact test, BH-adjusted p-value < 0.01; see **Supplementary Dataset 2**).

**Figure 4.**
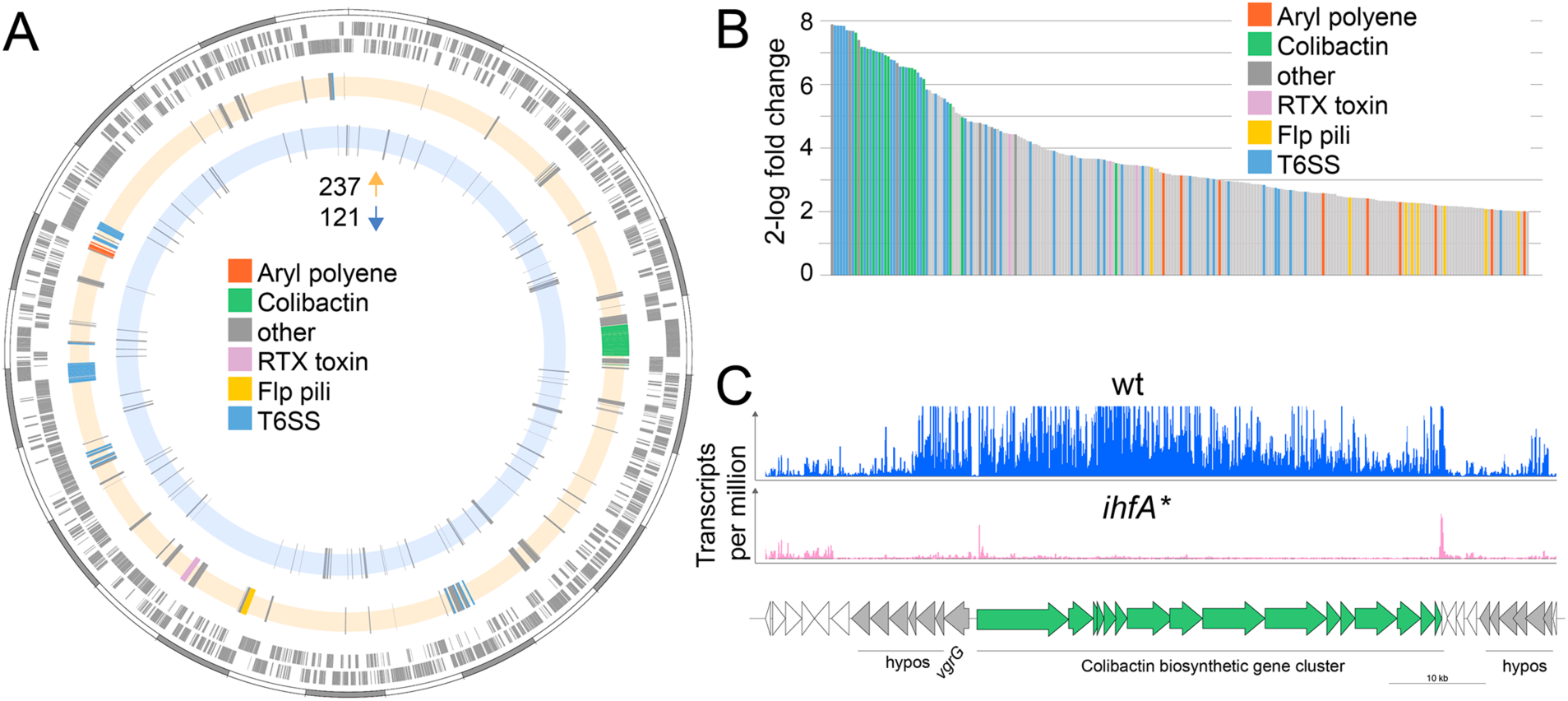
Differential gene expression between *F*. *perrara* wt and *ihfA*\* mutant during *in vitro* growth. (**A)** Chromosomal localization of all genes significantly differentially expressed (2-log fold change = |2|, Fisher’s exact test p value <0.05, FDR <0.05) between *F*. *perrara* wt and the *ihfA\** mutant. Starting from outside, the first circle shows the scale of the genome representation of *F*. *perrara* in gray and white steps of 100 kb. The second and third circles (gray) depict the genes on the plus and minus strands of *F*. *perrara*. The fourth (beige) and fifth (light blue) circle depicts genes upregulated and downregulated in wt compared to *ihfA\**. Genomic islands are highlighted by coloration. (**B)** Bar plot of the genes differentially expressed between *F*. *perrara* wt and *ihfA\** with a log2-fold change >2 (Fisher’s exact test p value <0.05, FDR <0.05). **(C)** Comparison of the transcriptional profile of the genomic location encoding the colibactin biosynthetic gene cluster between *F*. *perrara* wt and the *ihfA\** mutant. Transcripts per million were visualized using the Integrative Genome Browser (74). The colibactin operon is schematically depicted below (green arrows).

Genes belonging to these three categories include different subunits and effectors of the two T6SSs of *F. perrara*, the Clb biosynthesis gene cluster, various components of the Flp pili, and a RTX (repeats in toxin) toxin belonging to the Type I secretion system family (**Supplementary Dataset 1**). Also, the genes of the APE biosynthesis gene cluster were among the upregulated genes, which is in line with the production of the corresponding metabolites in the wt but not in the mutant strain (**Figure 2**). Interestingly, a relatively large proportion of the upregulated genes encoded hypothetical or poorly characterized proteins. In fact, genes without COG annotation were also enriched relative to the entire genome of *F. perrara* (Fisher’s exact test, BH-adjusted p-value < 0.01; see Supplementary **Dataset 2**). Many of the upregulated genes were organized in genomic islands, with the largest one including the biosynthesis gene cluster of Clb and many hypothetical protein-encoding genes (**Figure 4C**). T6SS and Clb biosynthesis genes were among the genes with the highest fold changes relative to *ihfA\** mutant (**Figure 4B**, 28 of 32 genes with log2-fold change >6). Moreover, 64% of the upregulated genes (152/237) belonged to the *F. perrara*-specific gene content as based on our previously published genome comparison of *F. perrara* PEB0191 with four other strains of the family Orbaceae (three of the genus *Gilliamella* and one of the genus *Orbus*, **Supplementary Dataset 1)** (37).

Among the 121 down-regulated genes, only COG category O (‘Posttranslational modification, protein turnover, chaperones’) was statistically enriched (Fisher’s exact test, BH-adjusted p-value < 0.01, see **Supplementary Dataset 2**). Moreover, only a small fraction (12%) belonged to the “*F. perrara*-specific genes”, and fewer genes were organized into genomic islands. A more detailed inspection of the annotation revealed that a large number of the down-regulated genes were involved in transport and metabolism (40 genes), transcriptional regulation (10 genes), and protein folding (8 genes), highlighting clear differences in the functional roles of the up- and down-regulated genes. The two genes with the highest fold change (log2-fold change <-5) both encoded transcriptional regulators. One of them, *dksA* (Fpe_01158), is located upstream of the *mrsA*/*mrsB* antioxidant system (Fpe_01159 to Fpe_01162), which was also among the down-regulated genes. The other one is part of the two-component regulator system *basS*/*basR* (Fpe_02097 and Fpe_02098), which has been reported to act as an iron- and zinc-sensing transcriptional repressor and activator in *E. coli* (47, 48). Taken together, these results show that many accessory genes known to be involved in symbiotic interactions (colibactin, Flp pili, T6SS) are upregulated in *F. perrara* wt as opposed to *ihfA*\*, providing a list of candidate genes responsible for the colonization defect of the *ihfA\** mutant.

### T6SS, pili, APE biosynthesis and Clb biosynthesis genes are expressed during bee gut colonization

To test if the genes upregulated *in vitro* in the wt relative to *ihfA\** were expressed *in vivo*, we determined the transcriptome of *F. perrara* wt at day 5 and day 10 post colonization. A total of 260 (149 up and 111 down) and 298 (162 up and 136 down) genes were differentially expressed at day 5 and day 10 post colonization relative to growth *in vitro* (log2-fold change >|2|, quasi-likelihood F-test with p<0.05 and FDR<5%, **Supplementary Dataset 3**). There was a considerable overlap of the differentially regulated genes between the two time points (115 and 80 shared up- and down-regulated genes, respectively). At both time points, the COG category ‘Carbohydrate transport and metabolism’ (COG G) was significantly enriched among the genes upregulated *in vivo* relative to the entire genome (**Supplementary Dataset 4**). In addition, at time point day 10, also the COG category (P) ‘Inorganic ion transport and metabolism’ was enriched (P adj < 0.01, Fisher’s exact test, **Supplementary Dataset 4**). Genes belonging to these two categories encoded transporters for different sugars (Phosphotransferase systems), iron, and transferrin (**Figure 5 – figure supplement 1, Supplementary Dataset 3**). In addition, a catalase gene and several genes for the biosynthesis of the amino acid tryptophan were upregulated at both time points. However, only 14 and 19 genes of those upregulated *in vitro* in the wt relative to the *ihfA\** mutant (see **Figure 5**), were also upregulated *in vivo* at day 5 and day 10 post-colonization, respectively (**Supplementary Dataset 3**). This was expected, because the *in vitro* RNAseq analysis had shown that these genes are already expressed in the wt when grown on mTYG agar, which we used as a reference condition for the *in vivo* analysis. Indeed, when comparing count-normalized gene expression (as measured by Transcripts per million (TPM)) across the different conditions, we found that most of the T6SS machinery, APE biosynthesis, pilus, and iron uptake genes were expressed at both time points *in vivo*, and to similar levels as *in vitro* (**Figure 5, Figure 5 – figure supplement 2**). Only the Clb genomic island and some of the VgrG-like T6SS effector genes had clearly lower TPM values *in vivo* than *in vitro*, yet higher than in *ihfA* in vitro* (**Figure 5C and 5F**). These results suggest that most of the genes upregulated *in vitro* in the wt relative to *ihfA\** are also expressed at high level by the wt *in vivo*.

**Figure 5.**
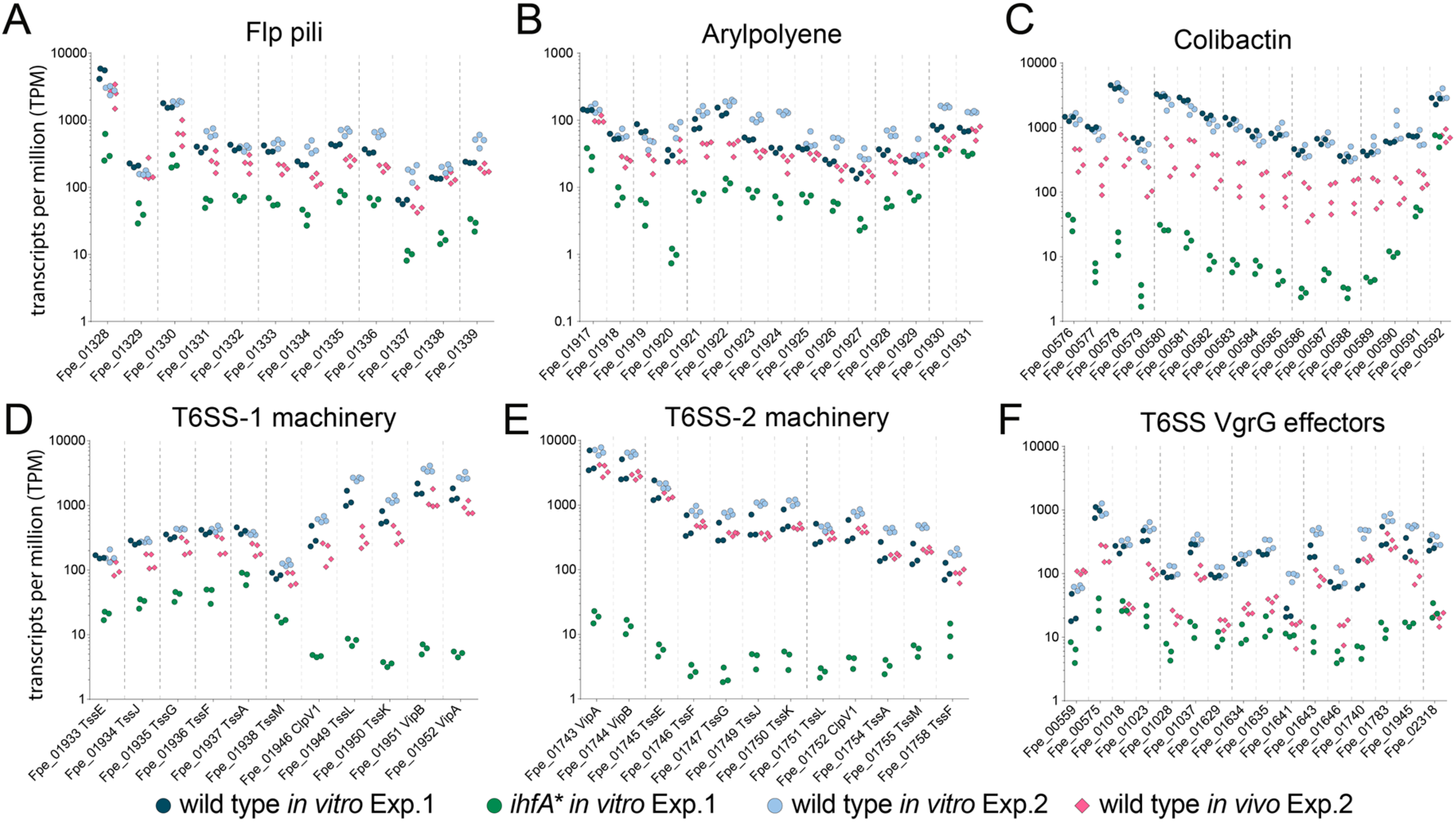
Gene expression of Ihf-regulated genes of *F. perrara* ten days post-inoculation of gnotobiotic honey bees. Transcripts per million were calculated for all replicates of the *in vitro* and the *in vivo* RNAseq experiments. For the in vitro experiment (Exp. 1), all three replicates of the wt and the *ihfA\** mutant are shown. For the *in vivo* experiment (Exp. 2), the four replicates of the day 10 time-point and the in vitro reference condition are shown. Data for day 5 time-point in comparison to day 10 time-point is shown in **Figure 5 – figure supplement 2**.

### Gene deletion of IhfA-regulated genes result in impaired gut colonization and/or abolish scab development

To test the direct impact of IhfA-regulated genes on host colonization and/or scab development we established a gene deletion strategy for *F. perrara* based on a two-step homologous recombination procedure (see methods and **Figure 6 – figure supplement 1**). This allowed us to create six different non-polar in-frame gene deletion mutants of potential colonization factors regulated by IHF (Supplementary Table S2). We deleted an essential gene of the colibactin biosynthesis pathway (Δ*clbB*), both *hcp* genes of the two T6SSs, either separately or as double mutant (Δ*hcp1,* Δ*hcp2,* and Δ*hcp1*/Δ*hcp2*), the gene encoding the major Flp pili subunit (Δ*pilE*), and the entire APE biosynthesis gene cluster (Δ*apeA-R*). Deletion mutants were confirmed by genome re-sequencing. The Δ*apeA-R* mutant was the only strain not forming yellow colonies anymore (**Figure 1 – figure supplement 1**), which is consistent with the idea that the aryl polyene pathway is responsible for the yellow color. We measured the growth of the gene deletion mutants *in vitro,* which was similar to the wt and *ihfA\** strains **(Figure 6 – figure supplement 2)**. Additionally, no significant differences were observed in cell length **(Figure 6 – figure supplement 3)**. Moreover, we corresponded OD_600_ to CFU counts and found that both the Δ*hcp1*/Δ*hcp2* and Δ*pilE* strains had lower counts than the wt strain **(Figure 6 – figure supplement 4)**. To compare the six gene deletion strains to *F. perrara* wt and the *ihfA\** mutant in terms of gut colonization and induction of the scab phenotype, each strain was inoculated into microbiota-free bees and CFUs assessed in the pylorus/ileum region ten days post inoculation. As expected, the wt successfully colonized the pylorus/ileum region of all analyzed gnotobiotic bees (median = 9.56*10^6^±5.06*10^6^ CFUs per gut; n=18 bees, two independent experiments) and induced the scab phenotype in 16 of 18 bees (**Figure 6A and Figure 6 – figure supplement 5**). In contrast, the *ihfA\** colonized poorly compared to the wt (median=1.46*10^4^±3.19*10^5^ CFUs per gut, n= 20 bees, Wilcoxon rank-sum test P<0.001), and not a single bee developed the scab phenotype. Of the six tested gene deletion mutants, only the mutant of T6SS-1 (Δ*hcp1*) reached wt colonization levels and also induced the scab phenotype in most of the bees (15 out of 20). The other five mutants all exhibited lower colonization levels than the wt strain. However, the severity of the colonization defect varied between the mutants, and while some of the mutants still caused the scab phenotype, others did not. For example, the Clb biosynthesis gene cluster mutant Δ*clbB* induced the scab phenotype in all but one bee, despite the fact that the colonization levels were about two-fold lower than for the wt (median=4.24*10^6^±2.87*10^6^ CFUs per gut). The gene deletion mutant of the T6SS-2 (Δ*hcp2*) and the double mutant of both T6SSs (Δ*hcp1*/Δ*hcp2*) reached similar colonization levels as the Δ*clbB* mutant. Yet, none of the bees colonized with these two mutants developed the scab phenotype, not even those that had very high bacterial counts (**Figure 6A**). The deletion mutant of the APE biosynthesis pathway (Δ*apeA-R*) showed again a different colonization phenotype: for some bees no colonization was detected, while others showed similar colonization levels as the wt of *F. perrara*. However, in contrast to the Δ*hcp2* and the Δ*hcp1*/Δ*hcp2* mutants, the Δ*apeA-R* mutant still induced the scab phenotype, but only in the bees that had high bacterial loads. Finally, the Δ*pilE* mutant had the strongest impact on colonization. Only five out of 20 bees had detectable levels of *F. perrara* in the gut (limit of detection: 75 CFUs per gut) and none of the bees developed the scab phenotype. Electron microscopy imaging revealed this strain did not have pili, confirming the Δ*pilE* mutation affects the formation of these structures **(Figure 6 – figure supplement 3C).**

**Figure 6.**
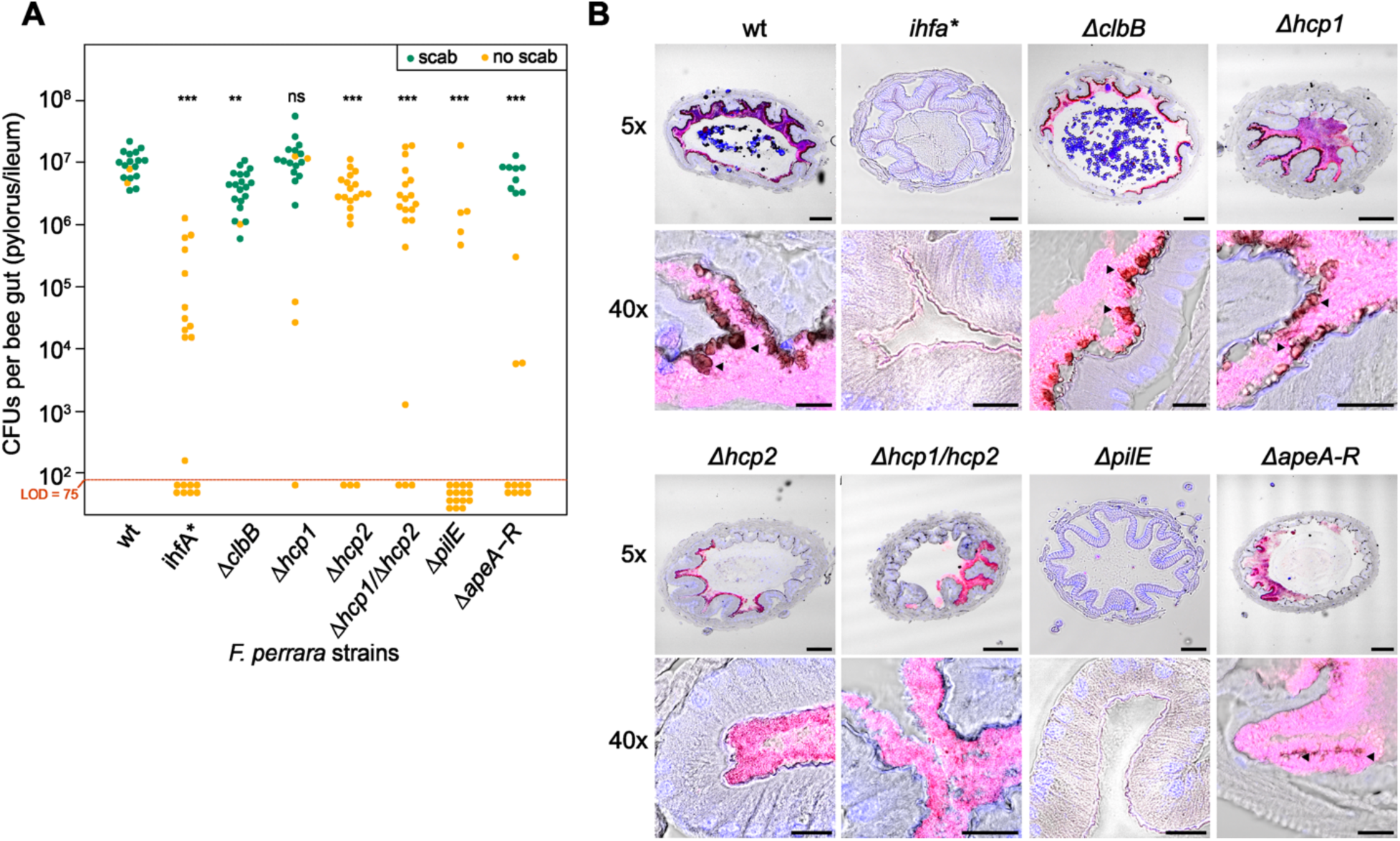
Gut colonization phenotypes of different gene deletion mutants of *F. perrara.* **(A)** Colonization levels were assessed 10 days after inoculation by counting colony forming units (CFUs) in dilutions of homogenized bee guts plated on BHI agar. Only the pylorus and ileum section of the gut were analyzed. Limit of detection (LOD) corresponds to the lowest colonization level detectable in our assay, i.e. points below the LOD correspond to bees for which no CFUs were detected. Statistically significant differences of the colonization levels of each mutant relative to the wt of *F. perrara* were determined using the Wilcoxon rank-sum test with BH correction. Bees were inoculated with an OD_600_ of 0.1. Data come from two independent experiments. **Supplementary Figure 15** shows the data points by experiments. * p<0.05, **p<0.01, ***p<0.001. Filled circle colors indicate whether a scab was detected during dissection (green = scab; yellow = no scab). **(B)** Location within the pylorus was assessed using FISH microscopy. Bees were inoculated with different *F. perrara* genotypes at OD_600_ =0.1, guts were dissected at day 10 after inoculation and sectioned using a microtome. Hybridizations were done with probes specific for *F. perrara* (magenta). DAPI counterstaining of host nuclei and bacteria is shown in blue. Images were generated by merging brightfield, *F. perrara* and DAPI images that were obtained for the same section of the gut. The composite images here shown were obtained by merging the images of each channel presented in **Figure 6 – figure supplements 6 and 7**. These were obtained using the 5x and 40x objectives of the Zeiss LSM900. Scale bar for images obtained with 5x: 100 µm, for 40x: 20 µm.

The Δ*clbB,* Δ*hcp2* and Δ*hcp1/*Δ*hcp2* mutants all reached similar colonization levels, yet only colonization with Δ*clbB* led to the development of the scab phenotype. These differences in scab formation could be due to an altered colonization pattern of the Δ*hcp2* and Δ*hcp1/*Δ*hcp2* mutants. To address this hypothesis, we visualized how the gene-deletion mutants are distributed in the pylorus (**Figure 6B, Figure 6 – figure supplements 6 and 7**). We obtained cross-sections of the pylorus of bees associated with these mutants, stained them with DAPI and a *F. perrara*-specific FISH probe and imaged these sections using confocal microscopy. As previously reported (34), *F. perrara* wt was found to colonize the pylorus region, forming a dense biofilm in close proximity to the host and occupying the crypts. A similar colonization pattern was observed for Δ*hcp1*, Δ*hcp2*, Δ*hcp1/*Δ*hcp2,* Δ*clbB* and Δ*apeA-R.* In contrast, for the *ihfA\** and Δ*pilE* mutants, we did not identify any bacteria in the analyzed gut sections which is in agreement with the low colonization levels of these mutants detected by CFU plating. Dark spots corresponding to scab material were found along the cuticular lining colonized by bacteria for the Δ*clbB,* Δ*hcp1* and Δ*apeA-R,* but not for the Δ*hcp2* and Δ*hcp1/*Δ*hcp2* mutants, which matches the visual inspections of the dissected guts for the presence/absence of the scab phenotype across the different strains (**Figure 6B**). Based on these results, we conclude that the inability of the Δ*hcp2* and Δ*hcp1/*Δ*hcp2* mutants to trigger the scab phenotype cannot be explained by an altered localization in the gut. Overall, these results confirm that IHF regulates various host colonization factors that when deleted cause distinctive colonization defects.

### *F. perrara* T6SS-2 and the APE biosynthesis pathway regulate gut colonization in complex bacterial communities

In natural conditions, *F. perrara* shares its niche with other symbionts of the bee gut. As several factors regulated by IHF are known for their role in microbial interactions (49, 50), we also wanted to test the impact of the six gene-deletion mutants on the ability of *F. perrara* to colonize the bee gut in the presence of other community members. We compared gut colonization levels of the gene deletion mutants, *ihfA*,* and the wt in the presence and absence of a synthetic community (BeeComm_002) composed of 13 strains representing major core microbiota members of the honey bee gut microbiota (detailed information in **Supplementary Table 4**). For five of the eight tested strains we did not detect a significant effect of the BeeComm_002 on the colonization levels (**Figure 7, Figure 7 – figure supplement 1**). This included *F. perrara* wt, which successfully colonized the gut independently of the presence of the BeeComm_002 (Wilcoxon rank-sum test P=0.899, two independent experiments), the *ihfA\** (Wilcoxon rank-sum test P=0.638) and the pili mutant Δ*pilE* (Wilcoxon rank-sum test P=0.217), which both already exhibited a strong colonization defect in mono-colonization, and the colibactin gene cluster mutant Δ*clbB* (Δ*clbB:* Wilcoxon rank-sum test P=0.127) and the T6SS-1 mutant Δ*hcp1* (Wilcoxon rank-sum test P=0.068), which both colonized at equal levels in both conditions (**Figure 7**). For the other three mutants we observed a significant reduction in colonization levels in the presence of the BeeComm_002. The colonization levels of T6SS-2 mutant Δ*hcp2* decreased seven-fold (median mono-association: 3.86*10^7^±4.16*10^7^, median BeeComm_002: 5.10*10^6^± 1.77*10^7^, Wilcoxon rank-sum test P=0.002) whereas both the T6SS double mutant Δ*hcp1/hcp2* (Wilcoxon rank-sum test P=0.029) and the APE biosynthesis mutant Δ*apeA-R* (Wilcoxon rank-sum test P=0.004) failed to colonize most bees tested **(Figure 7 and Figure 7 -figure supplement 1B)**. These results demonstrate that the T6SS-2 and the APE biosynthesis pathway play a role in regulating *F.perrara* colonization in the presence of other symbionts, possibly regulating the interaction with these bacterial partners.

**Figure 7.**
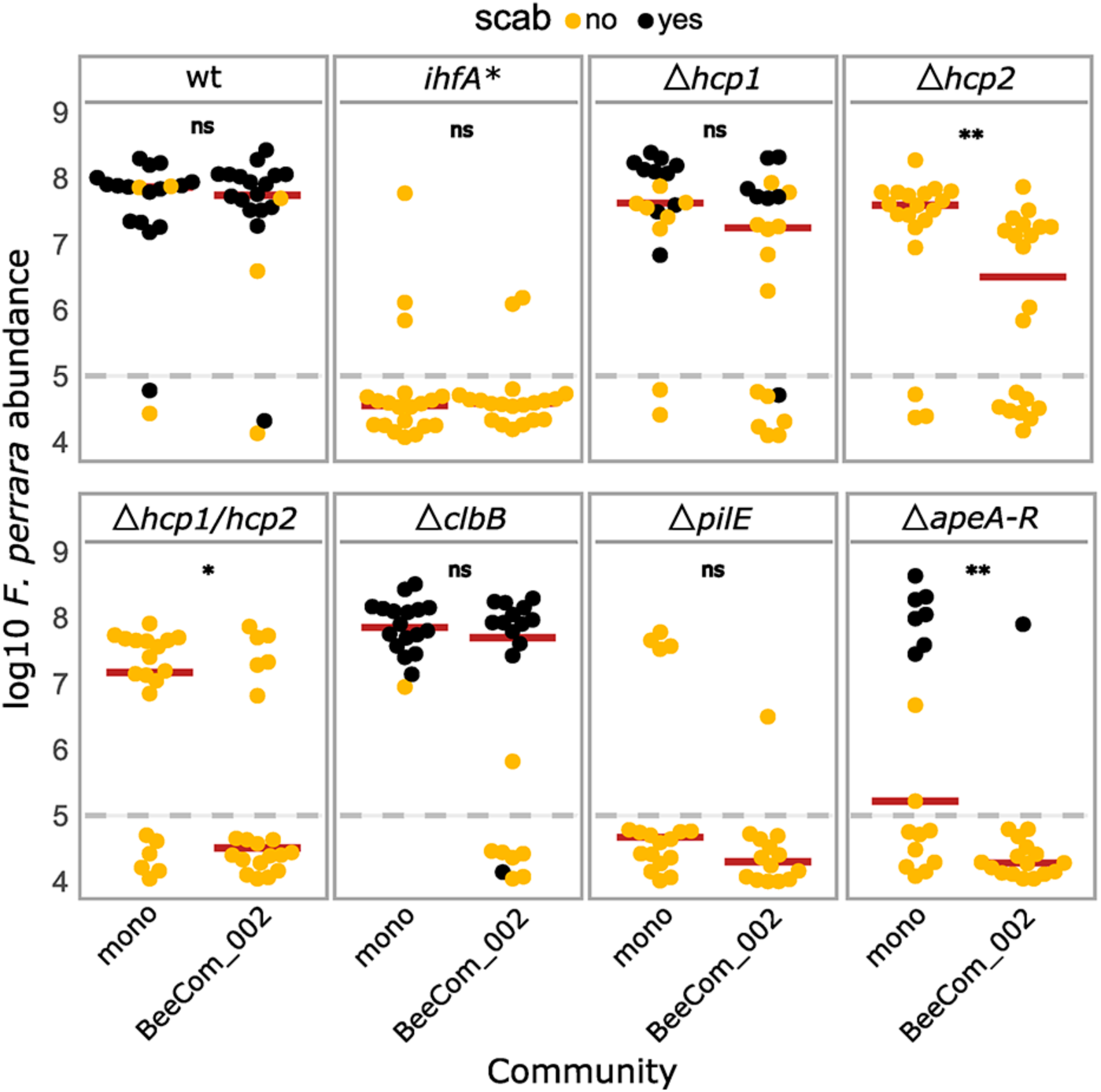
Gut colonization of the gene deletion mutants in the presence of bacterial competition. Bees were inoculated with *F. perrara* alone (mono) or in the presence of a defined bacterial community representing core members of the bee gut microbiota (BeeCom_002). Colonization levels were assessed 10 days after inoculation by qPCR. Only the pylorus and ileum section of the gut were analyzed. The dashed grey line refers to the limit of detection (LOD) and corresponds to the lowest colonization level detectable in our assay, i.e. points below the LOD correspond to bees for which no *F. perrara* was detected. Statistically significant differences between the colonization levels of each mutant in mono association compared to in the presence of the defined microbial community were determined using the Wilcoxon rank-sum test with BH correction. Data comes from two independent experiments. **Figure 7 – figure supplement 1A** shows the data points colored by experiments. * p<0.05, **p<0.01, ***p<0.001. Filled circle colors indicate whether a scab was detected during dissection (black = scab; yellow = no scab).

## Discussion

In this study, we identified genetic factors that allow the gut symbiont *F. perrara* to colonize its specific niche in the pylorus of the honey bee gut and to induce the scab phenotype. Our results advance the understanding of the genetic factors that facilitate symbionts to colonize niches in the animal gut. Specifically, we find that the DNA-binding protein IhfA plays an important role in gut colonization. IhfA is a histone-like, nucleoid-associated protein (NAP) that forms a heterodimer together with IhfB. IHF binds to and bends DNA in a sequence specific manner (40–42, 51), thereby facilitating the recombination of mobile DNA elements (40, 41) and influencing gene expression. We identified three different mutations in IHF, all located in the region of the protein interacting with the DNA. All three IHF mutants formed larger colonies than the wt strain on mTYG agar. Together, this suggests that the identified mutations change the DNA-binding properties of IHF resulting in broad transcriptional changes in *F. perrara* that provide a growth advantage *in vitro* relative to the wt strain.

*F. perrara* colonizes the epithelial surface of the pylorus, where it adheres to the cuticular lining and forms a thick biofilm-like layer (34). IHF has been shown to bind to extracellular DNA, which in the case of the human pathogen *Haemophilus influenzae* increases biofilm stability (52). Therefore, it is possible that IHF has a direct impact on gut colonization of *F. perrara* by stabilizing the biofilm formed on the host epithelium. However, based on our gene expression data and the results of the gene deletion experiments, it seems more likely that IHF influences gut colonization by regulating downstream genes involved in host interaction. Several of the genes regulated by IHF are known to play important roles in the adhesion to surfaces or the formation of biofilms. For example, pili are key factors for adhesion across many bacteria (53–56) and have already been shown in the case of the bee gut symbiont *S. alvi* to be beneficial for gut colonization and biofilm formation (7). In agreement with these previous results, several of the deletion mutants of IHF-regulated genes showed colonization defects in our gnotobiotic bee experiments corroborating that IHF impacts gut colonization through its effect on gene expression rather than through a direct role in biofilm formation.

The pili mutant (Δ*pilE*) had the strongest colonization defect of the tested gene deletion strains, likely because pili mediate the adhesion to the host epithelium, and therefore allow *F. perrara* to persist and replicate in the pylorus/ileum region of the honey bee gut. The deletion mutant of the APE biosynthesis pathway (Δ*apeA-R*) also showed a clear colonization defect. However, while some of the bees inoculated with this mutant had no detectable levels of *F. perrara* in the gut, others reached similar levels as bees colonized with the wt strain and also developed the scab phenotype. Aryl polyenes can protect bacteria from reactive oxygen species (46, 57) which are produced by insects as part of the host immune response in the midgut (58, 59). Our time course experiment revealed that in the first few days of colonization the bacterial numbers of *F. perrara* in the gut are much lower than in the initial inoculum (**Figure 2**) suggesting that the bacteria experience a significant population bottleneck at the beginning of the colonization process. It is possible that the Δ*apeA-R* mutant is impaired in its ability to resist reactive oxygen species (or other physicochemical stressors) when passing through the anterior sections of the honey bee gut and therefore reaches its niche in the pylorus in only a fraction of the inoculated bees. These stressors may be even more active in the presence of a complex community, which could explain the even lower colonization success of Δ*apeA-R* in the presence of the BeeComm_002. However, APEs have also been implicated in other functions such as biofilm formation (57). Therefore, future studies will be needed to elucidate how the APE biosynthetic pathway of *F. perrara* impacts honey bee gut colonization. Given the widespread occurrence of these biosynthetic genes throughout different bacteria, this might help to understand their role in host colonization in a much wider context (43).

A somewhat unexpected result of our mutant analysis was that the deletion of one of the genes encoding an essential NRPS/PKS enzyme (ClbB) of the Clb biosynthetic gene cluster did not affect scab development and had only a weak impact on gut colonization (2-fold lower levels than wt). Studies on *E. coli* have shown that the Clb biosynthesis pathway induces DNA damage in eukaryotic cells (60) and contributes to tumorigenesis in the mammalian gut (61, 62). We have previously shown that *F. perrara* also induces DNA damage in eukaryotic cells and that this is dependent on a functional Clb biosynthesis pathway (37). Therefore, we had speculated that the genotoxic activity of colibactin may trigger the local melanization response and the development of the scab phenotype upon colonization with *F. perrara* (17, 34). However, the results presented here show that colibactin is not causing the scab phenotype. Therefore, other characteristics of *F. perrara* must explain why this bacterium causes the scab phenotype.

Usually, bees that have <10^6^ bacteria in the gut do not develop visible scab at day 10 post colonization (**Figure 3 and Figure 7**), which suggests that a certain number of bacteria is needed to elicit this characteristic host response. On the contrary, not all bees with high levels of colonization developed a scab phenotype indicating that high loads are necessary but not sufficient to trigger the development of a visible scab phenotype. Specifically, the *Δhcp2* mutant of T6SS-2 and the *Δhcp1*/*Δhcp2* double mutant of T6SS-1 and T6SS-2 both reached relatively high colonization levels, yet not a single bee developed the scab phenotype. T6SS are usually involved in interbacterial warfare by injecting toxins into neighboring bacterial cells (63–65). However, there is an increasing number of studies showing that certain T6SS effectors can also target eukaryotic cells and modify diverse eukaryotic processes, including adhesion modification, stimulating internalization, cytoskeletal rearrangements and evasion of host innate immune responses (66). While *F. perrara* is not the only bee gut symbiont harboring T6SS, the effector protein repertoires differ tremendously between different species and even strains of the same species (67). Thus, it is possible that some of the effector proteins of *F. perrara* may target the host rather than other bacteria eliciting the melanization response and scab phenotype in the bee gut. Additionally, both the *Δhcp2* and the *Δhcp* double mutant had increased colonization defects in the presence of the BeeComm_002. This raises the hypothesis that the T6SS-2 may be important for *F.perrara* to interact with other gut symbionts, namely bacteria of the genera *Snodgrassella* and *Gilliamella* that also colonize the pylorus. It is important to mention that our experimental design favored *F. perrara* over the individual members of the BeeComm_002 as, in the inoculum fed to the bees, *F. perrara* was 13 times more abundant than any member of the defined community. It is likely that a stronger colonization defect would be observed if the proportions of *F. perrara* to other community members were more even. In any case, the presence of the BeeComm_002 led to a reduction of the number of bees colonized by most *F. perrara* strains, but not for the wt **(Figure 7 – figure supplement 1B)**. This is particularly interesting in the case of the *Δhcp1* and *ΔclbB* mutants, suggesting a possible role for these genes in the presence of other symbionts.

We did not carry out a genome-wide screen for host colonization factors, and only generated deletion mutants of a few of the IHF-regulated genes. Thus, there are probably many other factors that also contribute to gut colonization. For example, our *in vivo* RNA-seq experiment indicated that metabolic genes involved in tryptophan biosynthesis, sugar transport, and iron uptake are upregulated during gut colonization. This is in line with the TnSeq screen performed in *S. alvi,* which revealed that genes of essential amino acid biosynthesis pathways and iron uptake are important colonization factors of the honey bee gut (68). It is however remarkable that *F. perrara* only upregulates the tryptophan biosynthesis pathway, which suggest a specific demand for the production of this specific amino acid during gut colonization.

While our study revealed that IHF is important for regulating host colonization factors in the honey bee gut, it remains to be elucidated whether IHF is a direct regulator of the identified genes. In *Vibrio fluvialis*, IHF binding sites were identified upstream of several T6SS gene clusters, indicating that direct transcriptional regulation of similar genes by IHF exists in other bacteria (69). Alternatively, IHF may act upstream of another regulator that controls the expression of the identified genes. In the plant pathogen *Dickeya zeae,* IHF was suggested to positively regulate different virulence factors through binding to the promoter region of a diguanylate cyclase gene, increasing the production of the secondary messenger c-di-GMP (70). In *F. perrara,* we found that the transcriptional regulators BasR and DksA were substantially downregulated in the wt versus the *ihfA\** mutant. Hence, it is possible that at least some of the differentially regulated genes may not be under the direct control of IHF but regulated via BasR or DskA.

In conclusion, we identified important gut colonization factors of the bee gut symbiont *F. perrara* and show that they are regulated by IHF. The wide occurrence of these genes in host-associated bacteria suggests similar roles in other environments and calls for a more detailed functional characterization. Our approach to create clean gene deletion mutants expands the available genetic toolbox for bee symbionts (33) and will help to dissect the molecular mechanisms of the identified gut colonization factors in this animal model.

## Material and methods

### Bacterial cultivation

*F. perrara* strains were cultivated on modified tryptone yeast extract glucose (mTYG) medium (0.2% Bacto tryptone, 0.1% Bacto yeast extract, 2.2 mM D-glucose, 3.2 mM L-cysteine, 2.9 mM cellobiose, 5.8 mM vitamin K, 1.4 µM FeSO_4_, 72.1 µM CaCl_2_, 0.08 mM MgSO4, 4.8 mM NaHCO3, 1.36 mM NaCl, 1.8 µM Hematine in 0.2 mM Histidine, 1.25% Agar adjusted to pH 7.2 with potassium phosphate buffer), Columbia Blood Agar (CBA) containing 5% defibrinated sheep blood (Oxoid)) or Brain Heart Infusion broth (BHI) and incubated in anaerobic conditions at 34-35 °C (8% H_2_, 20% N_2_, 78% CO_2_ in a Vinyl Anaerobic Chamber, Coy Lab). Fresh bacterial cultures were used for each experiment. To this end, pre-cultures streaked out from glycerol stocks were grown for 48 hours and re-inoculated onto fresh plates for 16-24 hours of growth. For liquid cultures, pre-cultures streaked out from glycerol stocks were grown on TYG or BHIA plates for 48 hours and then inoculated into fresh liquid TYG or BHI. Cultures were incubated at 34-35 °C in a ThermoMixer C (Eppendorf) at 800 rpm for 16-24 hours of growth. Strains of *F. perrara* used in this study are listed in **Supplementary Table S2**.

### Rearing and experimental colonization of honey bees

Microbiota-depleted bees were generated as described by (18). For experimental mono-colonization, bees were starved for 1-3 hours by removal of the sugar water solution. Then, bees were cooled down to 4 °C in a refrigerator or on ice to transfer them (head side first) into 1.5 ml microfuge tubes with a hole at the bottom. Tubes with bees were kept at room temperature. For inoculation, each bee was fed 5 µl of *F. perrara* resuspended in sugar water:PBS (1:1 v/v) through the hole at the bottom of the microcentrifuge tube. The bacterial inocula were adjusted to an OD_600_ of 0.01 or 0.1 depending on the experiment. Colonized bees were kept at 30 °C with 70% humidity while having access to sugar water and sterilized bee pollen *ad libitum* until sampling.

### Tissue dissection and bacterial quantification from colonized honey bees

Bees were anesthetized by exposure to CO_2_. The whole gut or desired gut tissue, e.g., pylorus or ileum was dissected using a scalpel. Malpighian tubules were removed from the gut tissue and the presence of a scab was documented using a dissection stereomicroscope (Leica) as described in (14, 17). The tissues were placed into 2 ml screw-cap tubes containing glass beads (0.75-1 mm diameter, Roth) and 500-1000 µl PBS depending on the experiment. Homogenization of the sample was done by bead beating (FastPrep-24 5g MP Biomedicals) for 40 seconds at a speed of 7.5 m/s. Serial dilutions (1:10) were performed for each homogenate and plated onto mTYG or BHI agar. Single colonies were counted to determine the total number of bacteria per gut tissue by multiplying with the dilution factor.

### RNA sequencing of bacterial *in vitro* cultures

Fresh *F. perrara* cultures were prepared on mTYG plates, harvested, and directly transferred into a tube containing TRI reagent (Sigma-Aldrich, Merck) and silica beads (0.1 mm diameter, Roth). Samples were immediately snap frozen in liquid nitrogen and stored at −80 °C until RNA extraction. RNA was extracted using a modified TRI reagent protocol. Samples were homogenized by bead beating with a FastPrep instrument with CoolPrep adapter (FastPrep-24 5G, cooled with dry ice) for two cycles of 45 seconds at speed of 6 m/s, including a 30 seconds break between each cycle. Samples were kept at room temperature for 5 min, subsequently extracted using chloroform and RNA was precipitated using isopropanol overnight at −20 °C. Precipitated RNA was dissolved in RNase-free water (Gibco) and incubated with DNase (NEB) to degrade remaining DNA. RNA samples were cleaned up using NucleoSpin RNA clean up kit (Machery-Nagel) according to manufacturer’s protocol, eluted in RNAse-free water and stored at −80 °C until further use. RNA quality was assessed using Nanodrop, Qubit RNA HS kit (ThermoFisher Scientific) and Bioanalyzer instrument (Agilent). High quality RNA samples were sent to Lausanne Genomic Technology Facility (GTF, University of Lausanne) for RNA sequencing. Samples were depleted of 16S rRNA using RiboZero reagent (Illumina) and Truseq stranded-RNA Zero libraries were generated (Illumina) before sequencing on an Illumina HiSeq2500 generating single-end 100 bp reads.

### RNA sequencing of bacteria during honey bee gut colonization

Microbiota-depleted bees were colonized with freshly grown *F. perrara*. A fraction of the inoculum was directly transferred into 2 ml tube containing TRI reagent (Sigma-Aldrich, Merck) and silica beads (0.1 mm diameter, Roth). These samples were immediately snap-frozen in liquid nitrogen and stored at −80 °C until RNA extraction, which was carried out in the same way as described in the previous section. For the *in vivo* RNA samples, bees were colonized with *F. perrara* as described above and sampled at day 5 and day 10 post inoculation. Guts were removed from anesthetized bees, the pylorus and first part of the ileum was dissected, and the presence of the scab was recorded. For each sample, 10 pylorus and ileum sections from bees coming from the same cage were pooled in a 2 ml screw cap tube containing 750 μl of TRI reagent (Sigma-Aldrich, Merck), glass beads (0.75-1 mm diameter, Roth), and silica beads (0.1 mm diameter, Roth). Immediately after collection, samples were snap frozen in liquid nitrogen and stored at −80 °C until RNA extraction. RNA was extracted as described in the previous section. At each time point, four biological replicate samples were used for RNA sequencing, i.e., bees for one sample came from independent cages of gnotobiotic bees. The GTF at the University of Lausanne generated the libraries for the sequencing of the *in vivo* RNA samples. Poly-A depletion was performed in order to enrich for bacterial mRNA and Ribo-zero rRNA depletion was performed in order to remove prokaryotic and eukaryotic rRNAs. Then TruSeq stranded mRNA libraries (Illumina) were generated. The eight libraries were sequenced on an Illumina HiSeq 2500 to obtain single-end 125 bp reads. Libraries of the *in vitro* RNA samples were generated as described before. The four libraries corresponding to the *F. perrara* inocula used for colonizing the bees of the four replicates were sequenced on an Illumina MiniSeq at the Department of Fundamental Microbiology (125 bp single-end reads). Two *in vivo* samples (sample identifier D10_3M and D5_1M) sequenced on the HiSeq 2500 at the GTF were included in the MiniSeq run which confirmed that the two runs gave comparable results.

### Differential gene expression and gene enrichment analysis

Raw FASTQ files provided by the GTF containing all reads and corresponding tags indicating whether they were accepted or filtered out according to the CASAVA 1.82 pipeline (Illumina). Only reads tagged as accepted were kept for further analysis. FASTQC (http://www.bioinformatics.babraham.ac.uk/projects/fastqc/) was used to control the quality of the data, followed by Trimmomatic (Trimmomatic-0.35) to trim adapters. Filtered and trimmed reads were mapped onto the *F. perrara* genome using Bowtie (bowtie2-2.3.2). Mapped reads were quantified using HTseq (Version 0.7.2). Differential gene expression analysis was done with the Bioconductor package EdgeR (71) using R scripts. For the in vitro RNAseq analysis, negative binomial models were fitted to the data and quantile-adjusted conditional maximum likelihood (qCML) common and tagwise dispersion were estimated. The conditional distribution for the sum of counts in a group was used to calculate the significantly differentially expressed genes using an Exact test with a false discovery rate <5%. Only genes detected in all samples with at least 1 count per million were used for the analysis. For the in vivo RNAseq analysis, we followed the recommendations specified in the EdgeR user guide for generalized linear models. In short, read counts were normalized by trimmed mean of M-values (72) producing scaling factors used by EdgeR to determine effective library sizes. Then, negative binomial generalized linear models were fitted for each condition and quasi-likelihood F-tests for each defined contrast (i.e., pairwise comparison between conditions) was used to assess the significance of differentially expressed genes. As for the in vitro RNAseq analysis, only genes with mapped reads with at least 1 count per million in all replicates and conditions were used for the analysis. Genes were considered as differentially expressed upon fulfilling the following criteria: (i) log2 fold change ≥ 2 (ii) a p-value of <0.05 and (iii) a false discovery rate <5%. Transcripts per million (TPM) were calculated as follows (73):

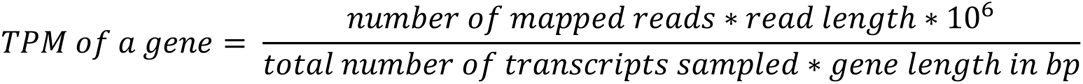

Coverage plots in **Figure 2A** and **Figure 4C** were generated with bam2wig.pl contained in the Bio-ToolBox-1.68 (https://github.com/tjparnell/biotoolbox) and visualized with the Integrated Genome Browser (74). Scripts for analyzing the RNAseq data can be found under the following Switchdrive link: https://drive.switch.ch/index.php/s/kCNTp4g7n60ffMi.

### DNA extraction of dissected gut tissue homogenates

DNA/RNA extraction was performed using an adapted hot phenol protocol from (18) consisting of sample thawing on ice, homogenization by bead beating at 7.5 m/s for 40 seconds (FastPrep-24 5g MP Biomedicals), two phenol extractions, and an ethanol precipitation overnight at −80 °C. DNA/RNA mixture was pelleted and eluted in RNase free water (Gibco). Samples were treated with RNaseA and DNA was purified using a Gel and PCR purification kit (Machery-Nagel). For those samples displayed in Figure 7 and Figure 7 – figure supplement 1, samples were homogenized with 165µL of a solution containing GI lysis buffer, Quiagen^TM^, and lysozyme (10:1 concentration) zirconia beads (0.1 mm dia. Zirconia/Silica beads; Carl Roth) and glass beads in a Fast-Prep24 5G homogenizer (MP Biomedicals) at 6 m/s for 45 s. After homogenization, samples were incubated at 37°C for thirty minutes. Then, 30µL of Proteinase K were added and samples were incubated at 56°C for one hour. The purification of nucleic acids was performed using CleanNGS magnetic beads (CNGS-0005) and the Opentron OT-2 pipetting robot. Purified DNA extracts were stored at −20°C until further use.

### Bacterial quantification by qPCR

Bacterial absolute abundances were determined using quantitative PCR (qPCR) assays targeting the 16S rRNA gene of *F. perrara*. Normalization was based on the number of host actin gene copies as described in (18, 75). Primer sequences and other primer characteristics are given in **Supplementary Table S3**. qPCR was conducted on a StepOne Plus instrument (Applied Biosystems) with the following run method: a holding stage consisting of 2 min at 50 °C followed by 2 min at 95 °C, 40 cycles of 15 sec at 95 °C, and 1 min at 65 °C. A melting curve was generated after each run (15 sec at 95 °C, 20 sec at 60 °C and increments of 0.3 °C until reaching 95 °C for 15 sec) and used to assess specificity of PCR products. qPCR reactions were performed in 10 μL reactions in triplicates in 96-well plates, and each reaction consisted of 1 μL of DNA, 0.2 µM of forward and reverse primer and 1x SYBR green “Select” master mix (Applied Biosystems). The qPCR reactions for the data corresponding to Figure 7 and Figure 7 – figure supplement 1 were carried out in 384-well plates on a QuantStudio™ 5 (Applied Biosystems). The thermal cycling conditions were as follows: denaturation stage at 50°C for 2 min followed by 95°C for 2 min, 40 amplification cycles at 95°C for 15 s, and 60°C for 1 min. Each reaction was performed in triplicate in a total volume of 10 µL (0.4 µM of each forward and reverse primer; 5 µM 1x SYBR® Select Master Mix, Applied Biosystems; 1 µL DNA). Each DNA sample was screened with two different sets of primers targeting either the actin gene of *A. mellifera*, or the universal 16S rRNA region.

For each target, standard curves were generated for absolute quantification using serial dilutions (from 10^7^ to 10 copies) of the target amplicon cloned into the plasmid vector. Absolute abundance of bacteria was calculated by using the standard curve and were normalized by the median actin copy number per condition (to account for differences in gut size) and by the amount of 16s rRNA copies per genome. For the calculation we used the following formula:

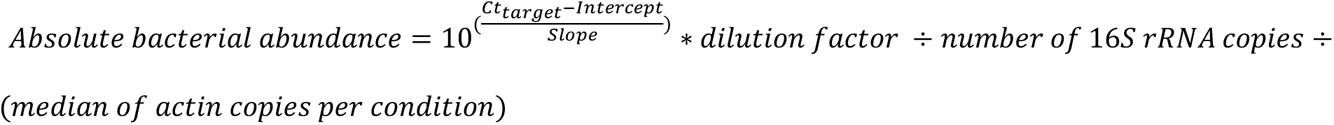

Here, Ct_target_ is the cycle threshold of the target bacterium and intercept and slope correspond to the values calculated for the standard curve of the target bacterium.

### Single cell microscopy

*F. perrara* was freshly grown from stock on plate. Subsequently, liquid overnight cultures starting at an OD_600_ 0.1 were inoculated in TYG for the experiment of the Figure 1 – figure supplement 2, and in BHI for the experiment of the Figure 6 – figure supplement 3. Liquid cultures were grown overnight 16-24 hours with shaking at 34-35 °C in a ThermoMixer C (Eppendorf) in anaerobic atmosphere. For Figure 1 – figure supplement 2, OD_600_ was measured and adjusted to 0.1 – 0.01 and cells were distributed onto small agar patches on a microscopy slide (Milan S.A., Menzel-Gläser). Cells were observed under 1000 times magnification using a ZEISS imager M1 microscope with phase contrast (PH3) condenser. Pictures were acquired randomly on the slides with VisiView software, 8-bit images were corrected with ImageJ and the contrast was set at min-value of 0 and max-value of 4000. A 10 μM scale bar was added with ImageJ (76). For the experiment displayed in Figure 6 – figure supplement 3, OD_600_ was adjusted to 0.1 and 10µl of bacterial solution were distributed onto small patches containing BHI and 1.5% UltraPure^TM^ Agarose from Invitrogen^TM^. Images were obtained using a Nikon ECLIPSE Ti Series inverted microscope coupled with a Hamamatsu C11440 22CU camera and a Nikon CFI Plan Apo Lambda 100X Oil objective (1000x final magnification). A 20 μM scale bar was added with ImageJ. For both experiments, the MicrobeJ plugin for ImageJ was used to measure cell size and area using the acquired pictures (77).

### Isolation and identification of aryl polyene compounds

*F. perrara* was streaked from the −80 °C glycerol stock onto mTYG plates. Plates were placed in an anaerobic tank with a pack of AnaeroGel 3.5 L (AN0035A Thermo Scientific) at 37 °C and incubated. Under aerobic sterile conditions, colonies were picked and 5 mL of mTYG medium were inoculated in a Hungate tube. After purging the liquid for 1 h with gas (83% N_2_, 10% CO_2_, 7% H_2_ v/v) bacteria were grown at 37 °C. After one day of growth, *F. perrara* wt and the *ihfA\** mutant were individually cultivated in Hungate tubes containing 30 mL mTYG for three days. Pellets were harvested, extracted with dichloromethane and methanol, then extracts were analyzed by HPLC-HESI-HRMS. The data revealed a strongly UV-Vis absorbent ion peak at *m/z* 323.1647 [M+H]^+^. To enrich this compound, 400 mL of mTYG were inoculated with 1 mL of pre-culture in a Hungate tube and purged as described above. After 5-10 days all but a few mL of medium were harvested and 400 mL of fresh mTYG was added under sterile aerobic conditions, purged with anaerobic gas and incubated at 37 °C. A total of 3 L of mTYG were used to obtain 6.53 g of cells, which were stored at −80 °C. Light exposure was minimized during all following steps of the aryl polyene enrichment procedure. A mixture of 340 mL dichloromethane and 170 mL of methanol was added to *F. perrara* cell pellets. After 2 h stirring at room temperature (RT), the suspension was filtered and 250 mL KOH (0.5 M) were added. After an additional 1 h of stirring at RT the pH was set to 6 using H_2_SO_4_ (1 M). The organic layer was washed twice with 500 mL water, once with 250 mL brine, dried over Na_2_SO_4_ and concentrated. The extract was separated by reverse-phase high performance liquid chromatography (RP-HPLC, Phenomenex Luna 5 μm C18, φ 21.2 x 250 mm, 15.0 mL/min, λ=420 nm) with water +0.1% formic acid (solvent A) and MeCN +0.1% formic acid (solvent B). Solvent compositions of 5% B for 4 min, a gradient to 95% B for 18 min, 95% B for 9 min and 5% B for 4 min were used. Fractions around 24 min exhibited strong UV absorption at 420 nm and were further purified by RP-HPLC (Phenomenex Luna 5μ Phenyl-Hexyl, φ 10 x 250 mm, 2.0 mL/min, λ=420 nm) with 80% B for 5 min and a gradient of 80% B to 100% B for 30 min. Peaks around 17 min were combined, concentrated and analyzed by NMR spectroscopy. HPLC-HESI-HRMS was performed on a Thermo Scientific Q Exactive mass spectrometer coupled to a Dionex Ultimate 3000 UPLC system. NMR spectra were recorded on a Bruker Avance III spectrometer equipped with a cold probe at 500 MHz and 600 MHz for ^1^H NMR and 125 MHz and 150 MHz for ^13^C NMR at 298 K. Chemical shifts were referenced to the solvent peaks of DMSO-*d_6_* at δ_H_ 2.50 ppm and δ_C_ 39.51 ppm.

### Structural characterization of the aryl polyene

^1^H NMR in conjunction with HSQC data suggested approximately 15 methines and aromatic protons, one methoxy group and one methyl group (**Figure 2 - figure supplement 3 - 7**). From the COSY spectrum, four methines of the conjugated double bond system, and two of the aromatic protons could be connected (**Figure 2 - figure supplement 7 and 8**). HMBC correlations from the methyl group to an aromatic carbon at δ_C_ 156 ppm placed it in *ortho*-position of a hydroxy group and next to a singlet aromatic proton. The singlet of the methoxy-group was connected to a carbonyl group by HMBC correlations (**Figure 2 - figure supplement 9**).

### Construction of gene deletion mutants

Targeted in-frame gene deletions of *F. perrara* were constructed using a two-step homologous recombination procedure (see **Figure 6 – figure supplement 1**). In a first step, suicide plasmids were constructed harboring two adjacent 700-800 bp-long homology regions matching the up- and downstream region of the gene or gene cluster to be deleted. To this end, the two homology regions were amplified using 2 Phanta Max Master Mix (Vazyme) and cloned into pKS2 using Golden Gate assembly. pKS2 is a derivative of pSEVA312 (78) which contains a sfGFP expression cassette flanked by two BsaI restriction sites in the multiple cloning site. All clonings were done in *E. coli* DH5a/λpir. To validate successful cloning of the two homology regions, PCR and Sanger sequencing was performed. The plasmids derived from pKS2 containing the homology regions were shuttled into the diaminopimelic acid (DAP)-auxotroph *E. coli* JKE201 and delivered to *F. perrara* wt via conjugation using bi-parental mating. To this end, *E. coli* JKE201 carrying the plasmid of interest was cultivated for 16-20 hours at 37 °C in LB with 500µM DAP (Sigma, LB-DAP) and 30 µg/ml Chloramphenicol. Sub-cultures were made by diluting the culture at a ratio of 1:20 into fresh medium and incubated until exponential growth was reached. A fresh culture of *F. perrara* was prepared by growing it for 36 h on mTYG agar. *F. perrara* and *E. coli* were harvested in PBS and the OD_600_ adjusted to 10. Equal quantities of both bacteria were mixed and spotted onto a cellulose acetate membrane filter (0.2 µm, 25 mm, Huberlab, Sartorius) placed on BHI agar with 500 µM DAP, followed by incubation at 35 °C for 6 hours under microaerophilic condition. Filters with bacterial mixtures were removed from the plates with a forceps, placed into a tube with PBS. Bacteria were removed from the filter by pipetting, vortexing and agitation for 10 min at 1500 rpm (Thermomixer C, Eppendorf). The bacterial suspension was centrifuged at 8000x g for 10 minutes, the supernatant discarded to remove dead bacteria and the bacterial pellet was washed once with 1000 µl PBS before resuspending in a reduced volume, i.e. 100 µl PBS. The bacterial mixture was plated onto CBA and mTYG with selection antibiotic and incubated at 35 °C for 5 days under anaerobic condition. Colonies were picked after 4-5 days of incubation and expanded to fresh plates. DNA was isolated and five different PCRs performed to check for successful integration of the plasmid into the chromosomal region of *F. perrara* targeted for deletion. Positive clones (loop-in strains) were stocked in liquid mTYG containing 20% (v/v) glycerol at −80 °C until further usage. To select for bacteria that have lost the integrated plasmid, *F. perrara* loop-in strains were grown on mTYG agar and subsequently transformed with plasmid pYE1. pYE1 is a derivative of pBZ485 (79) containing the restriction enzyme *I-SceI* under the control of the IPTG-inducible *lac* promoter. The activity of the restriction enzyme *I-SceI* will negatively select bacteria containing pKS2-derivatives as two *I-SceI* sites are flanking the multiple cloning sites on pKS2. To generate electrocompetent *F. perrara* cells, the bacteria were cultivated from glycerol stocks on mTYG agar and incubated at 35 °C for 2 days under anaerobic condition. Bacteria from plate were inoculated in BHI broth with a starting OD_600_ of 0.1 and grown anaerobically at 40 °C for 12-16 hours. Subsequently, growth was stopped by placing cultures onto ice for 15 minutes. Cells were made electrocompetent by washing twice with ice-cold MOPS solution supplemented with 20% glycerol, with decreasing volumes per wash. Electrocompetent cells were mixed with >500 ng plasmid DNA and incubated for 15 minutes on ice before transferring into an electroporation cuvette. Electroporation was performed applying a voltage of 2.5 kV with resistance of 200 Ω and capacitance of 25 µF. Bacteria were immediately resuspended in BHI and incubated anaerobically for 6 hours at 35 °C to allow phenotypic expression. Bacteria were plated on CBA, mTYG and/or BHIA plates with 25 µg/ml Kanamycin and 100 µM IPTG to select for pYE1 and induce the restriction enzyme *I-SceI*. Plates were incubated in anaerobic atmosphere at 35 °C. After 3 to 5 days of incubation, bacterial colonies were replica plated onto fresh plates containing 15 µg/ml chloramphenicol (selection marker of pKS2-derivatives) or 25 µg/ml kanamycin (selection marker of pYE1) to identify clones that cannot grow on chloramphenicol anymore as a consequence of the loss of the integrated pKS2-derivative. DNA was extracted and a PCR screen performed with primers amplifying the gene targeted for deletion as well as the overspanning region of the deletion. All clones possessing the gene deletion were subsequently expanded on mTYG agar and stocked in glycerol as described above

### Correspondence between OD and CFU

For the wt, *ihfA*\* and the six gene-deletion mutants, bacteria were grown in BHIA plates for 3 days at 35°C in anaerobic conditions. On the day of the experiment, bacteria were harvested and a bacterial solution at OD_600_ =0.1 was prepared per each genotype. Serial dilutions were performed in a 96-well plate by transferring 10µl of OD_600_ =0.1 into 90µl of BHI medium followed by several 1:10 dilutions in BHI. For each genotype, 5 µl of all dilutions were plated on BHIA plates, incubated for 3 days at 35°C in anaerobic conditions and quantified to calculate the number of CFUs present in 5 µl of the initial OD_600_ =0.1 solution. The CFU values present in 5 µl of bacterial solution at OD_600_ =0.1 is of relevance as it corresponds to the volume fed to bees in the colonization experiments. Three experimental replicates were performed.

### Growth curves in liquid cultures

*F. perrara* was freshly grown from stock on BHIA plates. Subsequently, liquid overnight cultures starting at an OD_600_ =0.1 were inoculated in BHI and grown for 16-24 hours anaerobically with shaking at 35 °C and 600 rpm in a ThermoMixer C (Eppendorf) in anaerobic atmosphere. To obtain the growth curves, one 96-well plate was inoculated with different *F. perrara* strains. Each well contained 200µl of a bacterial solution at OD_600_ =0.05. Per strain, four wells were inoculated. Absorbance values were measured at 600nm every 20 minutes for 72 hours, using a BioTek^TM^ Epoch2 microplate reader. The plate was incubated in anaerobic conditions at 34°C and continuous orbital shaking for the duration of the experiment.

### FISH microscopy

Tissue sections and FISH experiments were performed as previously described (34). Briefly, the pylorus and ileum of gnotobiotic bees were dissected and fixed for five days in Carnoy’s solution (ethanol-chloroform-acetic acid, 6:3:1 [vol/vol]). Fixed tissue samples were washed three times for 1 hour in absolute ethanol, then incubated three times for 20 min in xylene, and finally infiltrated with paraffin three times for 1 hour at 60°C. Samples were placed into molds containing melted paraffin and then hardened by placing them into an ice slurry. Paraffin-embedded tissues were cut into serial 5-µm sections with a microtome (Leica), placed on coated microscopy slides, and cleared from paraffin with xylene. Sections were then hybridized overnight with fluorochrome-labeled oligonucleotide probes targeting the 16S rRNA of *F. perrara* and DAPI. Samples were imaged using the Zeiss LSM900 confocal microscope. Per bacterial genotype, one gut was processed and imaged. The *F. perrara* probe used is named PE1_TYE563_G2, has the sequence CCGCTCCAGCTCGCACCTTCGCT and a Cy3 flurophore.

### Electron Microscopy

The wt, *ihfa\** and *ΔpilE* strains were grown in BHIA plates for 3 days at 34°C in anaerobic conditions. The day before image acquisition, bacteria were harvested from the plates, transferred to liquid BHI at an initial OD_600_ =0.1 and were grown for 16 hours in anaerobic conditions with shaking. On the day images were acquired, bacteria were diluted 50 times. Each bacteria suspension was adsorbed on a glow-discharged copper 400 mesh grid coated with carbon (EMS, Hatfield, PA, US) during one minute at room temperature. Posteriorly, the meshes were washed with three drops of distilled water and stained for one minute with uranyl acetate (Sigma, St Louis, MO, US) at a concentration of 1%. The excess of uranyl acetate was drained on blotting paper and the grid was dried during ten minutes before image acquisition. Micrographs were taken with a transmission electron microscope Philips CM100 (Thermo Fisher Scientific, Hillsboro, USA) at an acceleration voltage of 80kV with a TVIPS TemCam-F416 digital camera (TVIPS GmbH, Gauting, Germany).

### Generation of the BeeComm_002 bacterial community

Thirteen bacterial isolates that represent the most abundant species found in the bee gut were grown individually and assembled together to generate a bacterial community named **BeeComm_002** (**Supplementary Table 4**). Each individual member was grown in agar plates of its respective preferential medium and culture conditions for 3 days, bacteria were harvested from the plates, diluted to OD_600_ =1 in PBS, mixed together in the same proportions and placed in a glycerol stock (20% final concentration) at −80°C. *Gilliamella* strains were grown in BHIA plates and incubated in anaerobic conditions at 35°C. *Bifidobacterium*, *Lactobacillus* and *Commensalibacter* strains grew anaerobically and at 35°C in Man, Rogosa and Sharpe agar plates supplemented with fructose and L-cysteine. *Snodgrassella alvi* was grown in Trypticase Soy agar plates, *Bartonella apis* grown in Columbia Broth agar plates supplemented with 5% sheep blood and both isolates were placed in micro-aerophilic conditions with 5% CO2 and at 34°C.

### Co-colonization of bees with *F. perrara* and BeeComm_002

Each gene-deletion mutant, the wt and the *ihfA\** mutant of *F. perrara* were grown as previously mentioned and diluted to OD_600_ =1 on the day the colonization experiment was performed. 100 µl of a given *F. perrara* genotype at OD_600_ =1, 100 µl of BeeComm_002 at OD_600_ =1 and 800 µl of a PBS and sugar water solution (1:1) were mixed together. As BeeComm_002 is composed of 13 isolates and was mixed in a 1 to 1 ratio with *F. perrara*, *F. perrara* was 13 times more abundant than any individual member of this synthetic community. For the control treatments where BeeComm_002 was not added (mono-associations), 100 µl of a given *F. perrara* genotype at OD_600_ =1 were diluted in 900 µl of a PBS and sugar water solution (1:1). Gnotobiotic bees were generated, manipulated and inoculated with bacteria as previously mentioned. Ileum/pylorus regions were dissected at day 10 after inoculation and scab formation was assessed before snap-freezing the samples in liquid nitrogen. Samples were placed at −80°C until DNA was extracted using previously mentioned methods. Bacterial abundance was quantified using the approach described in the Methods section “Bacterial quantification by qPCR**”**. Primers used in this experiment are found in **Supplementary Table S3.**

### Genome re-sequencing

All gene deletion strains and the three white variants of *F. perrara* PEB091 were subjected to Illumina genome re-sequencing at Novogene (2×150bp). Mutations in the re-sequenced genomes relative to the genome of the wt strain of *F. perrara* were analyzed with *breseq* (80). The detailed results of this analysis can be found: https://drive.switch.ch/index.php/s/kCNTp4g7n60ffMi.

### Statistical analysis

Statistical analysis was performed using GraphPad Prism v6.01 or R. Results of all statistical tests can be found in **Supplementary Dataset 5**. Normality was analyzed using D’Agostino-Pearson and/or Shapiro-Wilk test. For comparison with two conditions, t-test was used for parametric data and Wilcoxon sum-rank test or Kolgomorov-Smirnov test for non-parametric data. For comparison of the growth curves of *F. perrara* wt and *ihfA\**, we used a permutation test ‘compareGrowthCruves’ included in the statmod package of R (81). For the analysis of the CFU counts, a linear model with binomial distribution was calculated with CFUs as the dependent variable and Genotype and Experimental Replicate as independent variables. Post-hoc comparisons between genotypes were performed using the emmeans R package (https://cran.r-project.org/web/packages/emmeans/index.html). For the comparison of the cell lengths, Wilcoxon sum-rank test was performed.

## Data availability

RNA sequence datasets are available under NCBI Gene Expression Omnibus ID GSE189728. Code, scripts and numeric data files of experimental data have been deposited on Switchdrive: https://drive.switch.ch/index.php/s/kCNTp4g7n60ffMi.

## Acknowledgments

We thank Stephan Gruber for input regarding the characterization of the ihfA* mutant strain and general feedback for the manuscript. We are grateful to Nicolas Neuschwander and Lucie Kesner for helping with some of the bee experiments. We thank Tania Trabajo for her help with the single cell imaging. This project was funded by the ERC-StG ‘MicroBeeOme’ (grant agreement No. 714804), the Swiss National Science Foundation (grant agreement No. 31003A_160345 and 31003A_179487), the NCCR Microbiomes (all awarded to PE). J.P. acknowledges funding by the European Research Council (ERC) under the European Union’s Horizon 2020 Research and Innovation Program (grant agreement No. 742739).

## Figure Supplements

**Figure 1 – figure supplement 1.**
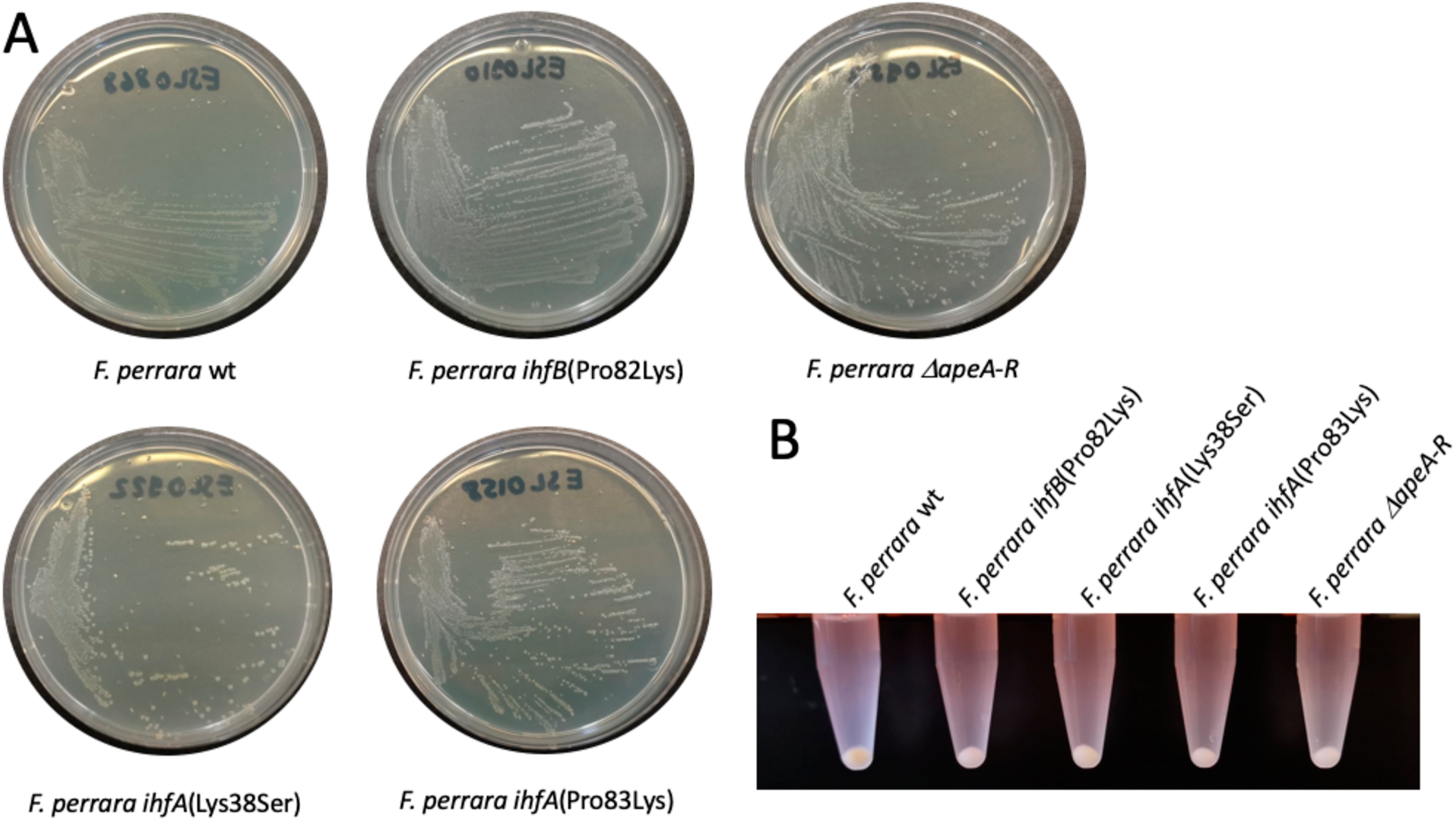
Colony morphology of different *F. perrara* strains on mTYG agar plates. **(A)** Dilution streaks of five different strains including the wt, the three separate point mutation strains of IHF (two in *ihfA* and one in *ihfB*), and the *ΔapeA-R* mutant after 2 days of growth under anaerobic conditions. **(B)** Bacterial cell pellets of the same strains as in **(A)** harvested from the agar plates, resuspended in 1x PBS, and centrifuged for 5 min at 5000 rpm.

**Figure 1 – figure supplement 2.**
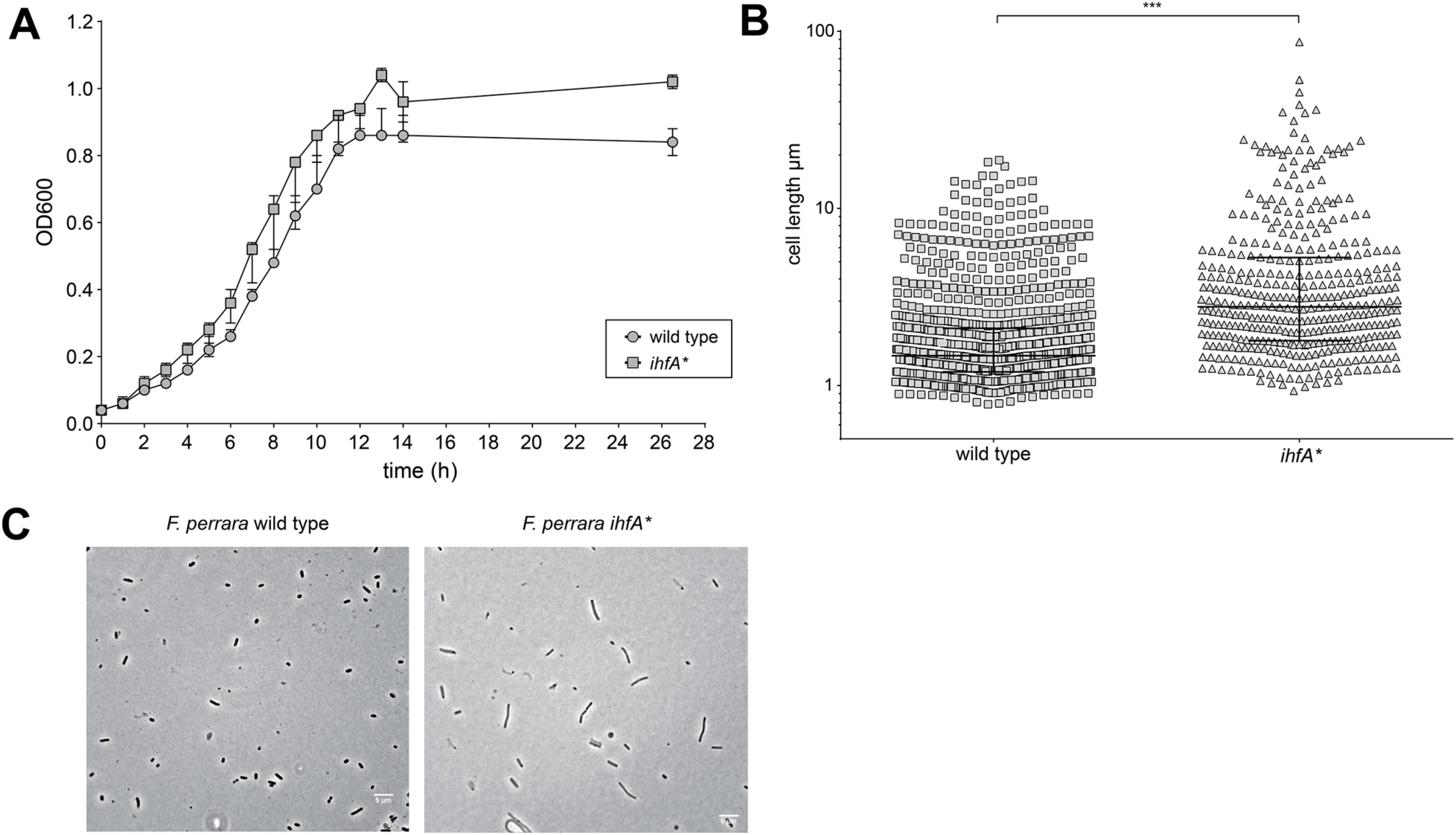
*In vitro* characterization of *F. perrara ihfA\**. **(A)** Growth curve of *F. perrara* PEB0191 wt and *ihfA\** in liquid TYG. The optical density (OD600) was measured every hour for 14 hours (Permutation test, p = 0.097). (**B)** Quantification of single cell length of *F. perrara* wt PEB0191 and *ihfA\** from the single cell microscopy experiment shown in C. Significance was tested using Kolgomorov-Smirnov test, stars depict significance: *** p<0.0001. **(C)** Single cell light microscopy of *F. perrara* wt and *ihfA\**. The scale bar (5µm) is depicted in the right lower corner.

**Figure 2 – figure supplement 1.**
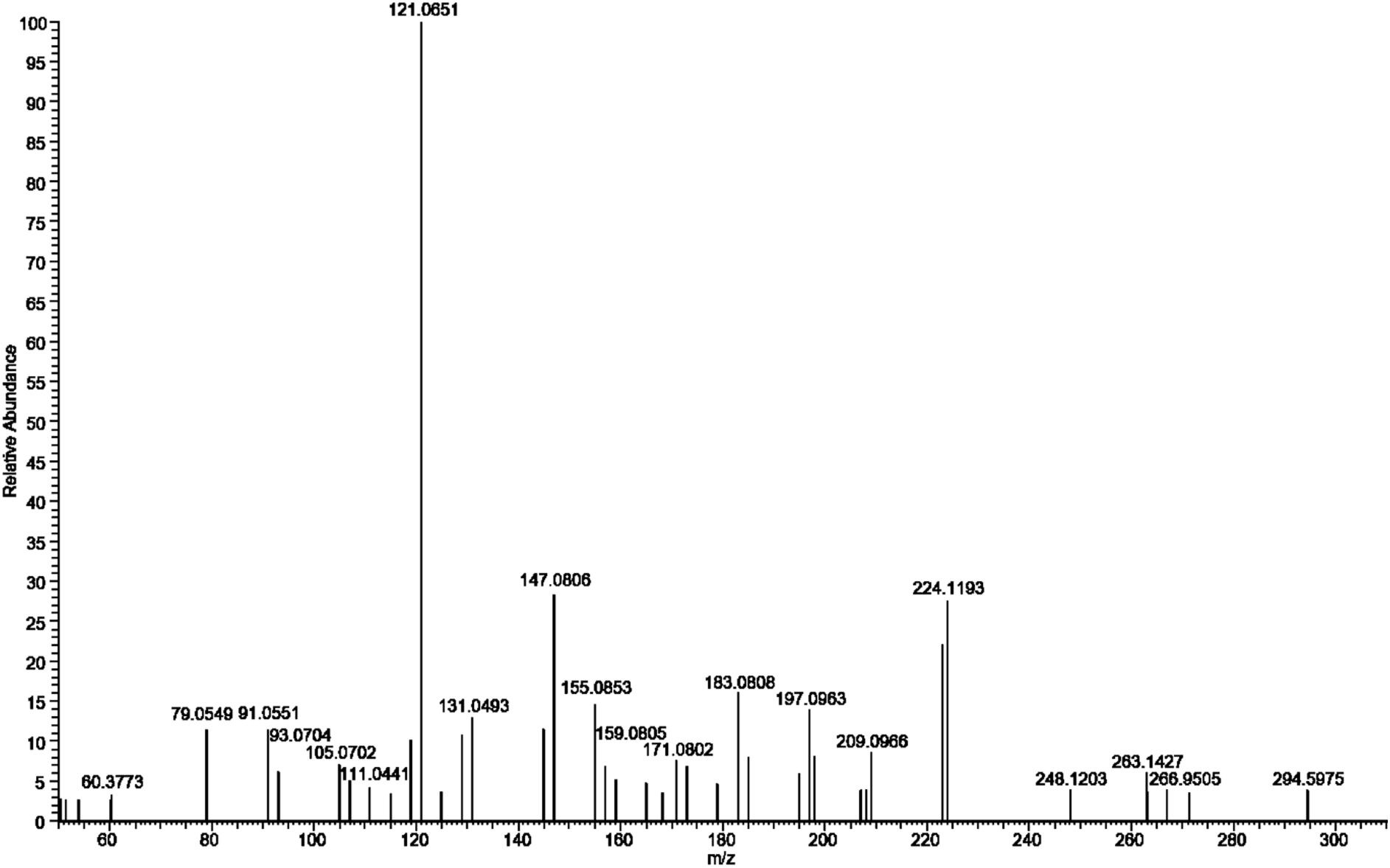
Ms-Ms fragmentation spectrum of *m/z* = 323.1647 in extracts of *F. perrara* wt. Many of the prominent ions were reported from aryl polyenes by (**46**): *m/z*= 121.00, 131.00, 145.00, 147.08, 171.08, 183.00, 197.08, 209.08, 223.08. The spectrum was recorded at 30 kV collision energy.

**Figure 2 – figure supplement 2.**
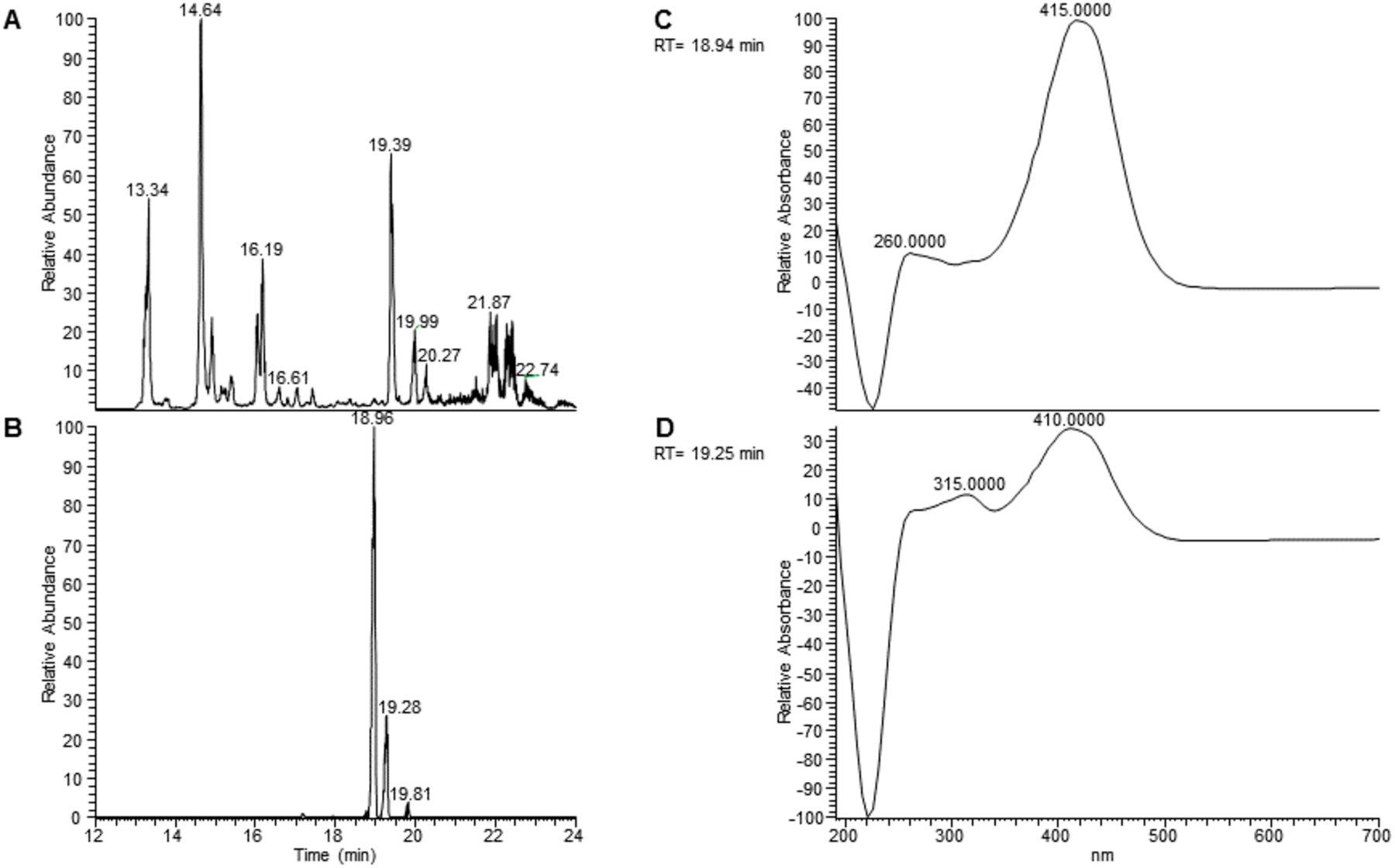
UV-spectrum of the extracts indicates isomerization of the aryl polyene. UV spectrum of the extracts indicates isomerization of the aryl polyene. (**A)** Total ion chromatogram of extracts of *F*. *perrara* wt. (**B)** Extracted ion chromatogram of *m/z* = 323.1640 Da (+/-5ppm). Storage led to the formation of an additional peak at 19.28 min, possibly by isomerization. **C** The UV-spectrum at 18.94 min has the maximum around 415 nm, similar to that reported by (43). **D** The UV-spectrum at 19.25 min shows an additional maximum at 315 nm, characteristic for a *trans*-*cis* conversion of a double bond.

**Figure 2 – figure supplement 3.**
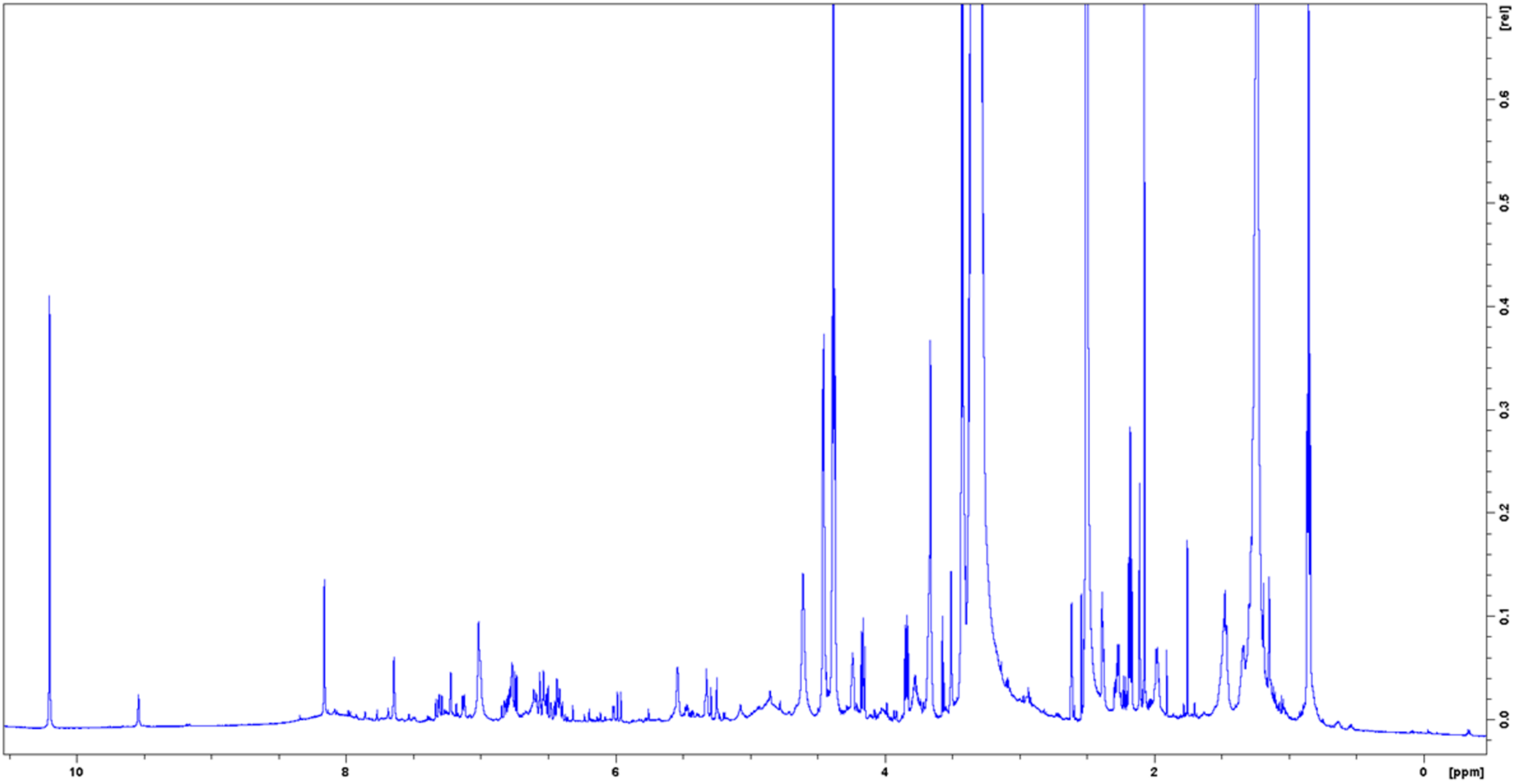
^1^H NMR spectrum of enriched aryl polyene in DMSO–δ_6_ at 298

**Figure 2 – figure supplement 4.**
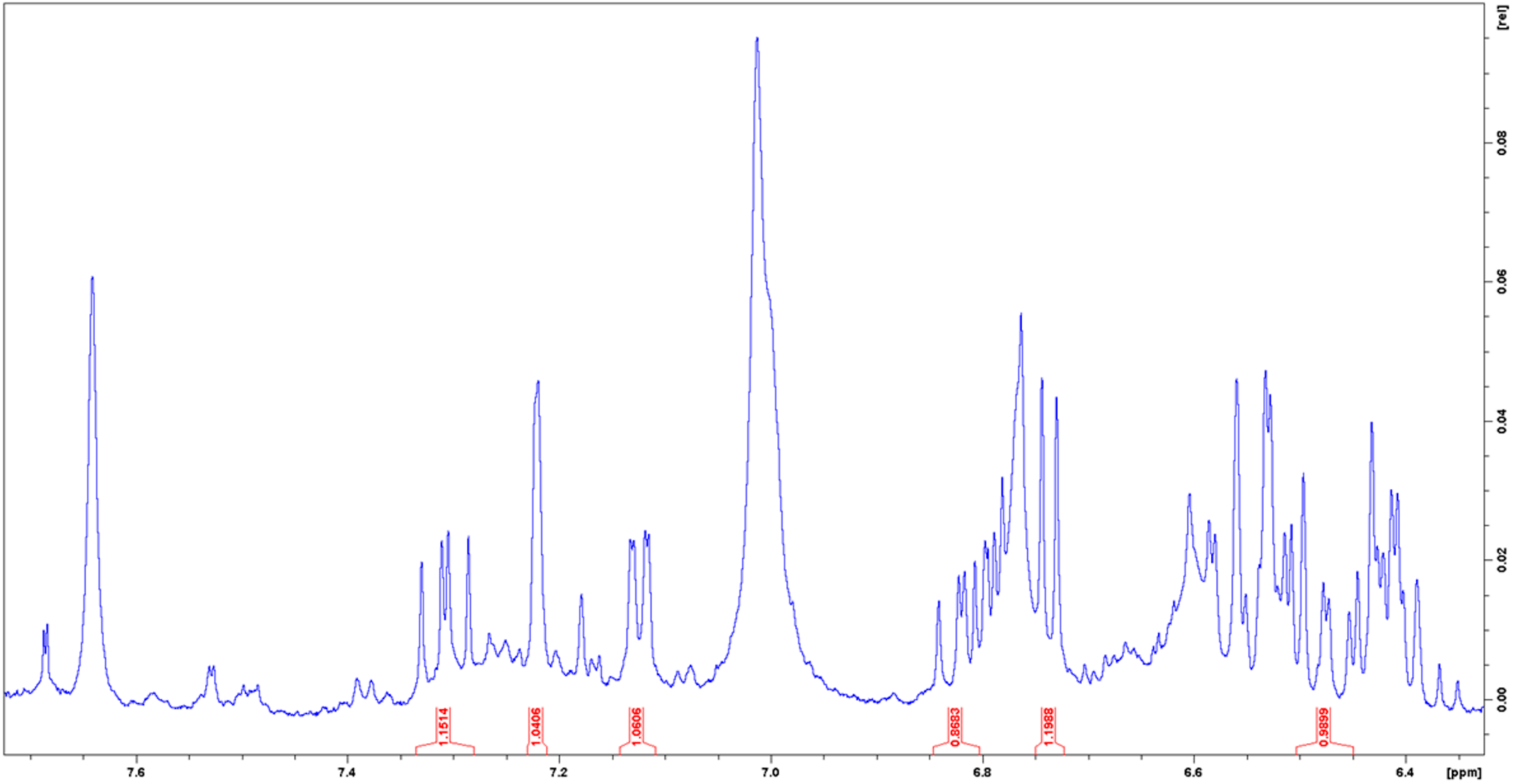
^1^H NMR spectrum of enriched aryl polyene in DMSO–*δ*_6_ at 298 K. Enhanced view of the aromatic region and integrals of assigned peaks.

**Figure 2 – figure supplement 5.**
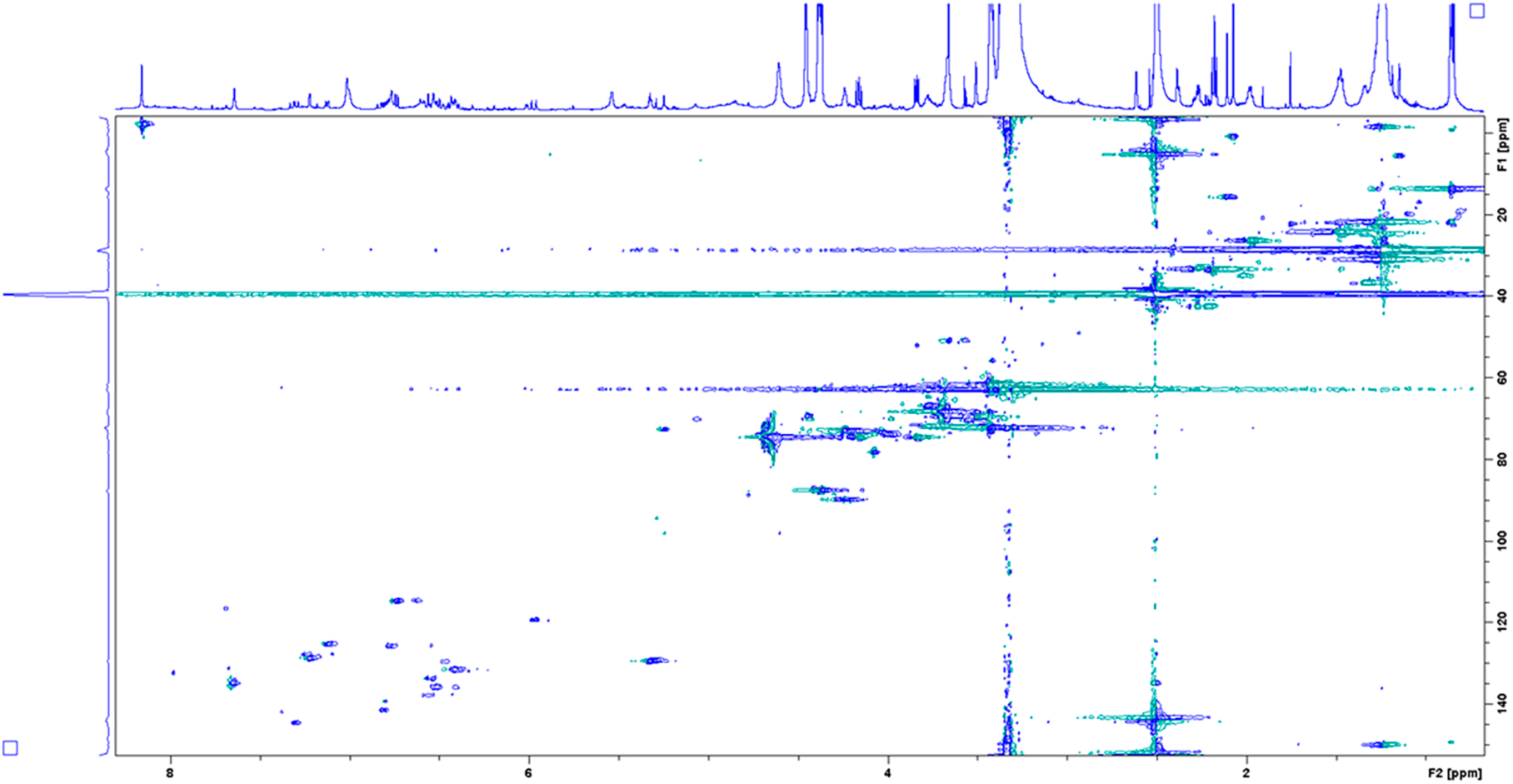
HSQC spectrum of enriched aryl polyene in DMSO–*δ*_6_ at 298 K.

**Figure 2 – figure supplement 6.**
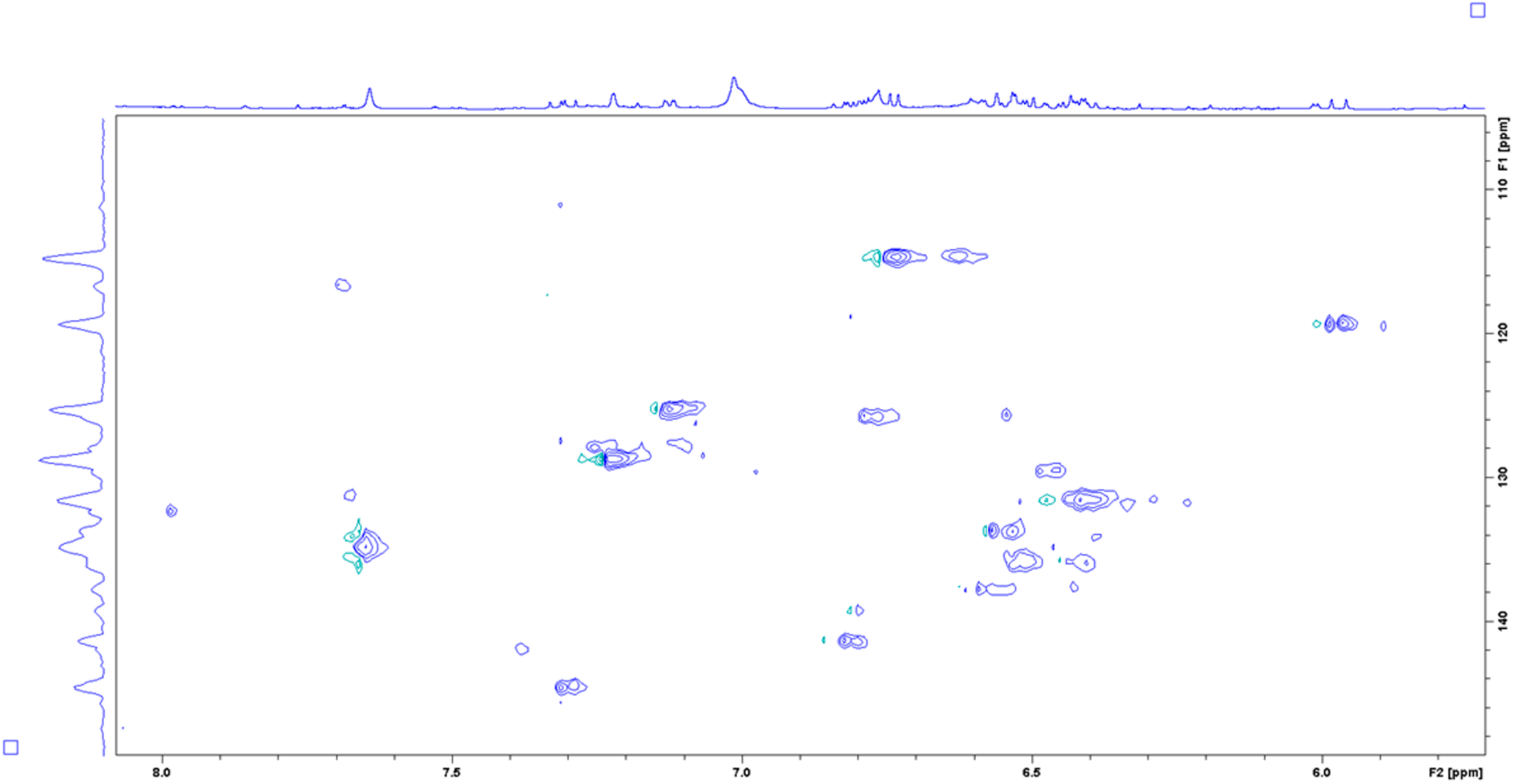
HSQC spectrum of enriched aryl polyene in DMSO–*δ*_6_ at 298 K. Enhanced view of the aromatic region.

**Figure 2 – figure supplement 7.**
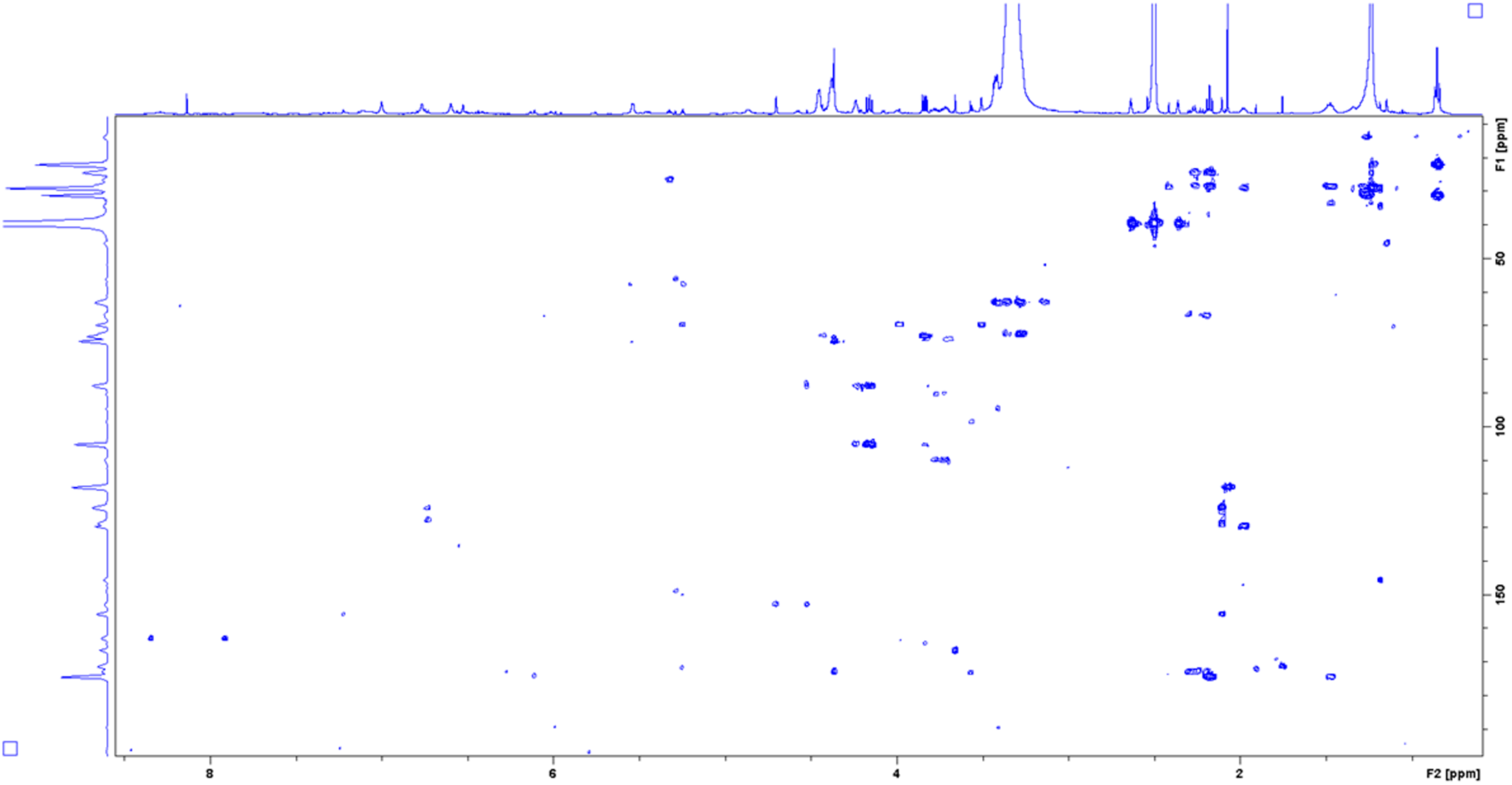
COSY spectrum of enriched aryl polyene in DMSO–*δ*_6_ at 298 K.

**Figure 2 – figure supplement 8.**
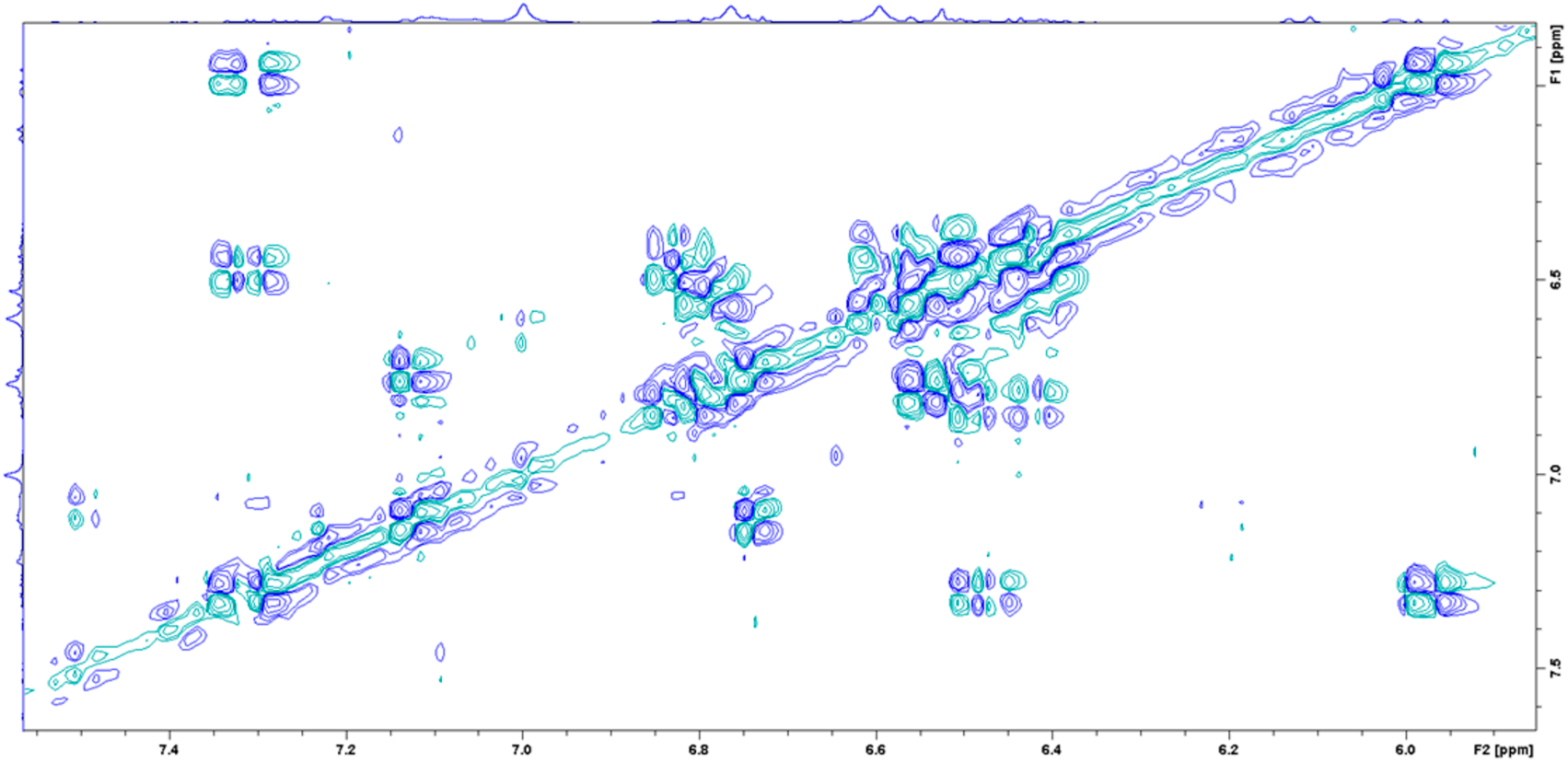
COSY spectrum of enriched aryl polyene in DMSO–δ6 at 298 K. Enhanced view of the aromatic region.

**Figure 2 – figure supplement 9.**
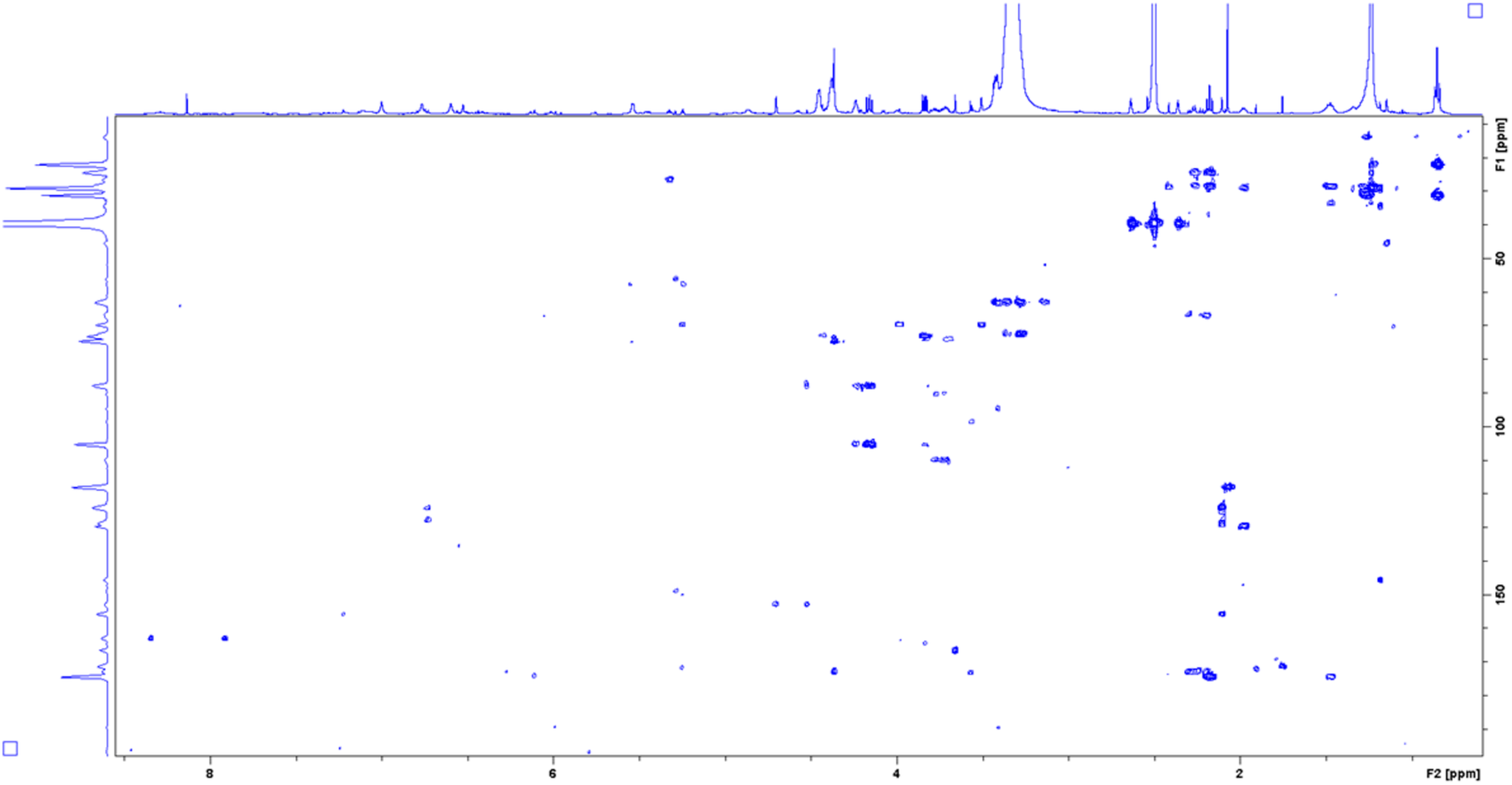
HMBC spectrum of enriched aryl polyene in DMSO–δ6 at 298

**Figure 2 – figure supplement 10.**
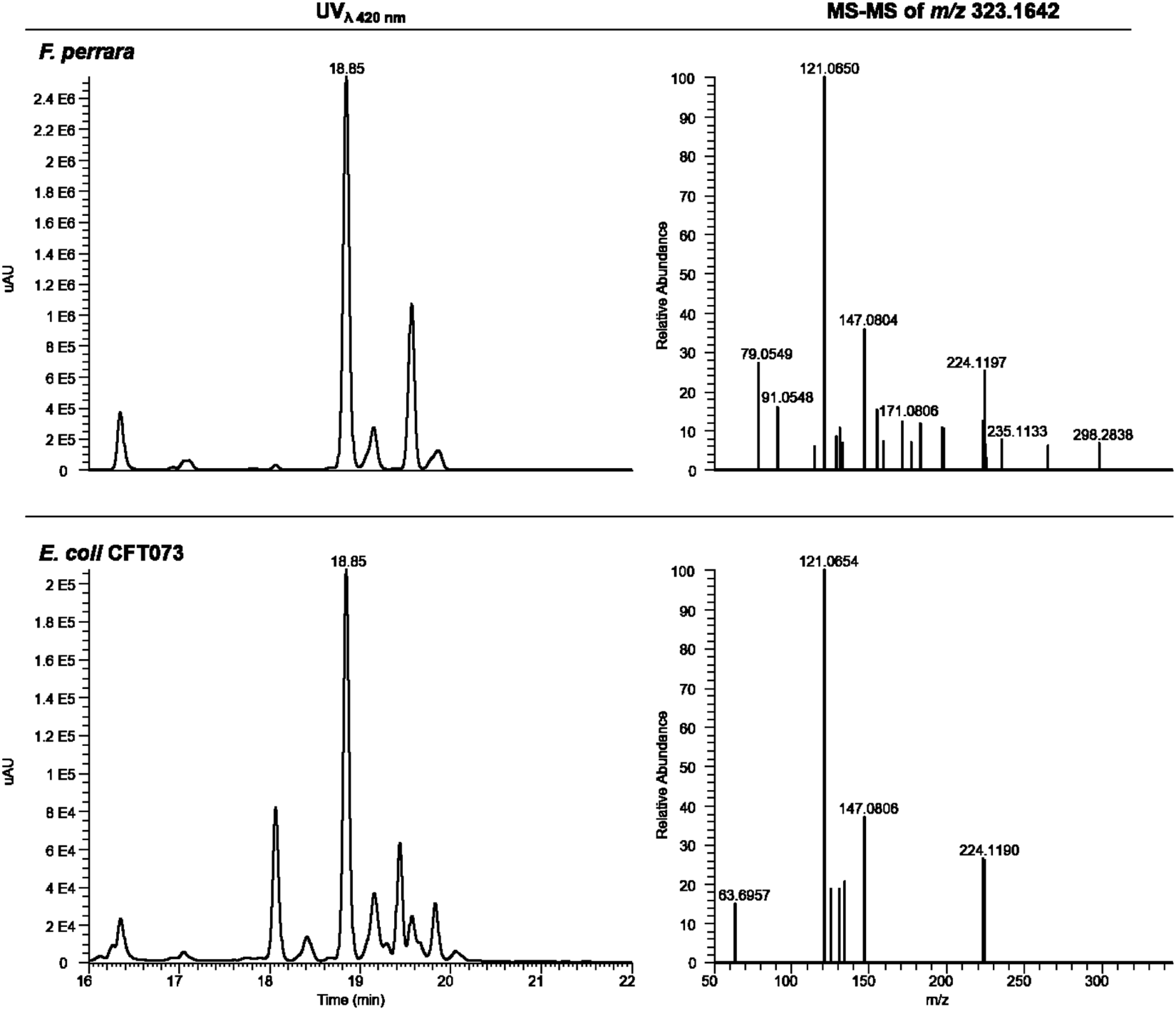
HPLC-HRESIMS analysis of *F. perrara* and *E. coli* CFT073. At 18.85 min both extracts show strong absorbance at λ= 420 nm. The MS-MS fragmentation pattern of the ion *m/z* 323.1642 at that retention time shows a similar pattern in extracts from both strains.

**Figure 3 – figure supplement 1.**
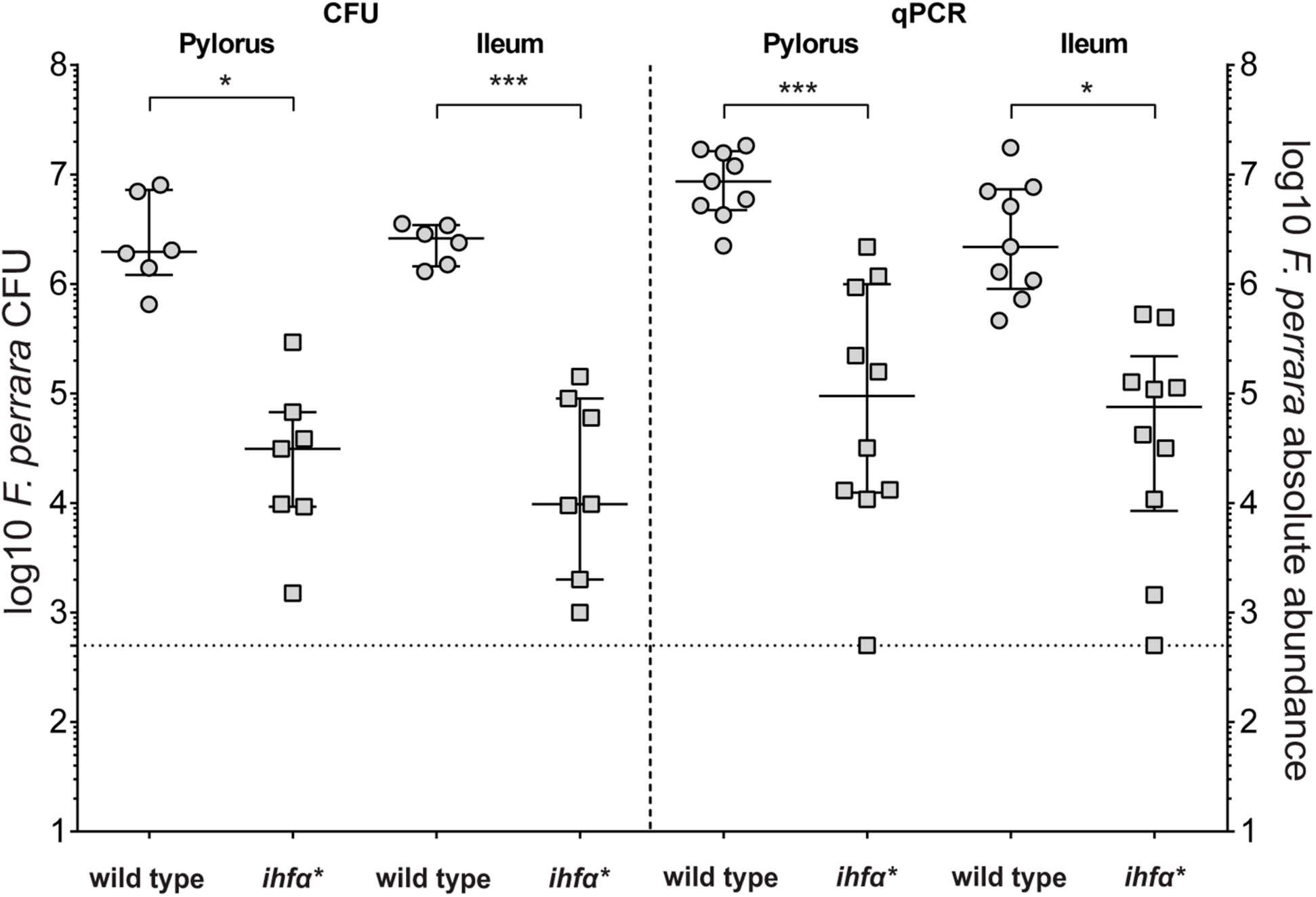
Quantification of *F. perrara* wt and *ihfA\** in pylorus and ileum. Mono-colonization of bees with *F. perrara* wt and *ihfA\**. Colonization levels are measured by colony forming units (CFU) and quantitative PCR (qPCR) 10 days post colonization in the specified gut regions: pylorus and ileum. Wilcoxon rank-sum test was used to assess the statistical significance of differences. n=6.

**Figure 5 – figure supplement 1.**
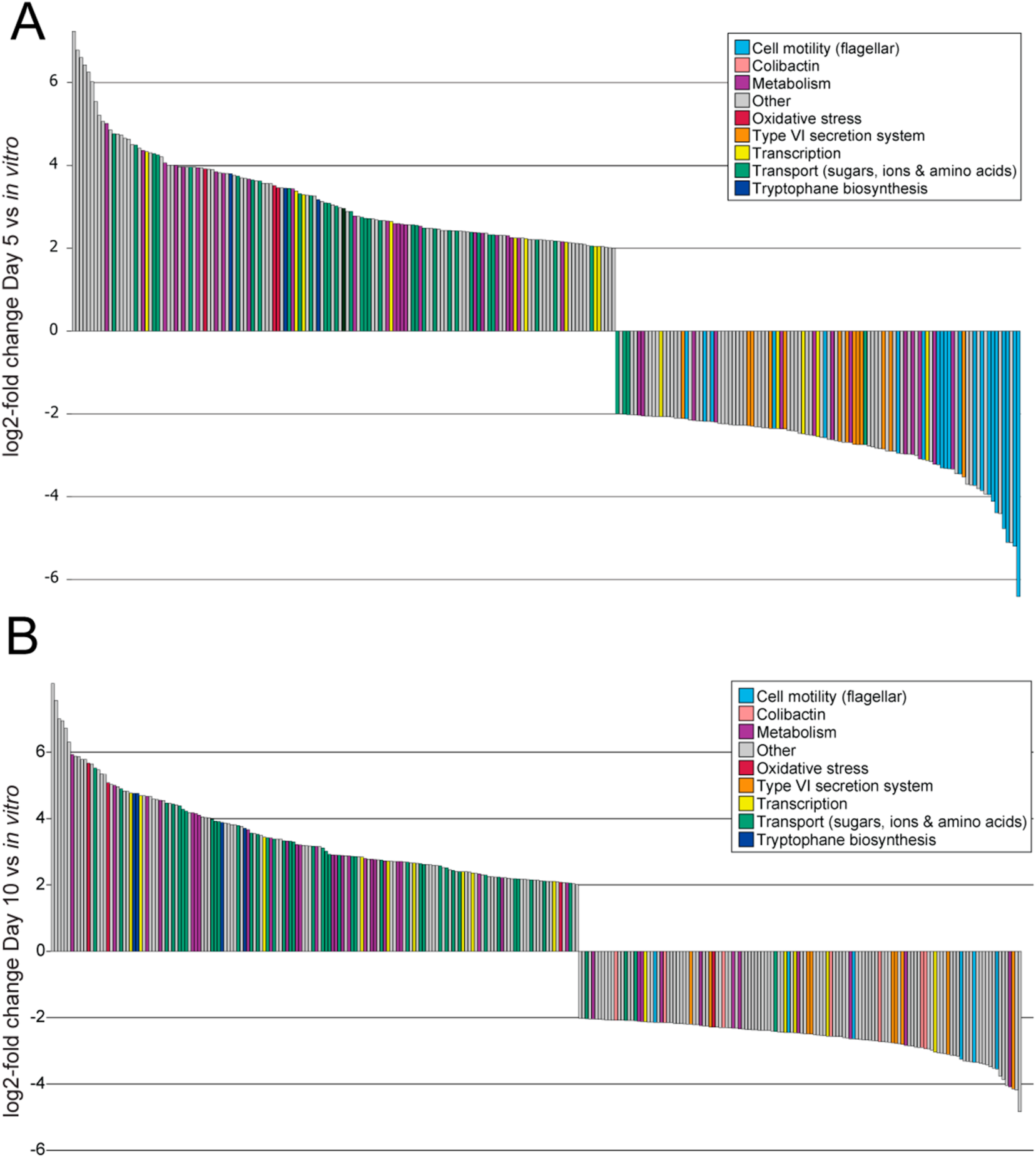
RNAseq comparison of *F. perrara* during *in vivo* colonization compared to growth *in vitro*. Significantly differentially expressed genes with a |log2-fold change| >|2| of *F. perrara* PEB0191 wt at day 5 (**A**) and day 10 (**B**) post colonization in comparison to growth *in vitro*. Genes are colored according to their category. The experiment was performed in quadruplicates, all genes displayed were significantly differentially expressed with p<0.05 and a FDR <0.05 (Exact test).

**Figure 5 – figure supplement 2.**
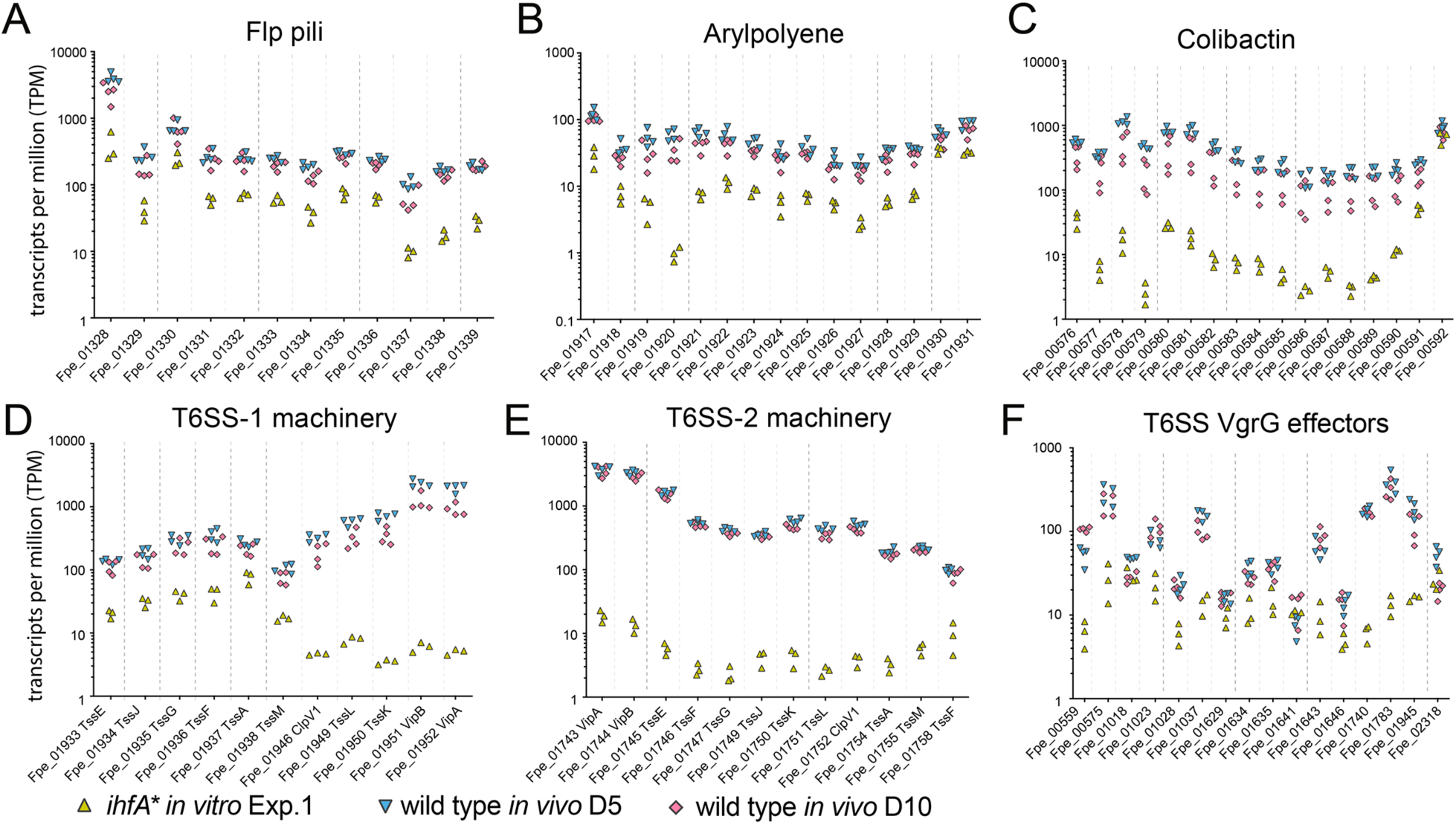
Gene expression of Ihf-regulated genes of *F. perrara* five days post-inoculation of gnotobiotic honey bees. Same as Figure 5, but only time point Day 5 and Day 10 are depicted in comparison to *F. perrara ihfA* in vitro*.

**Figure 6 – figure supplement 1.**
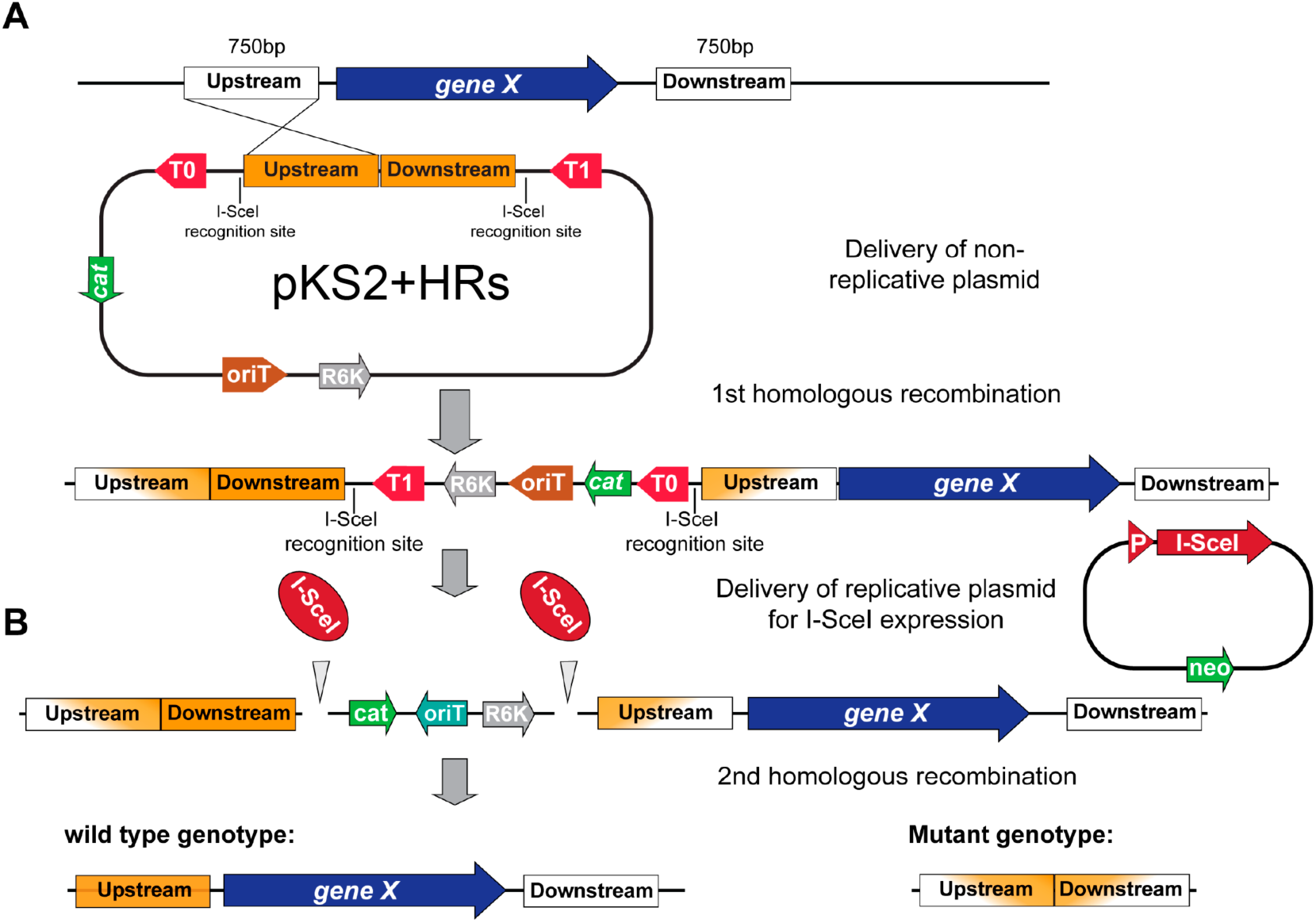
Scheme of the gene deletion strategy based on a two-step homologous recombination procedure. (**A)** A non-replicative plasmid pKS2 integrates via homologous recombination of one of the two cloned ‘homology regions’ (HRs) upstream or downstream of the gene that is targeted for deletion. (**B**) In a second step, a replicative plasmid harboring the restriction enzyme I-SceI is transformed. I-SceI targets corresponding recognition sites located on pKS2 resulting in the selection of either revertant (wt genotypes) or mutant genotypes that underwent a second homologous recombination event in the region targeted for deletion. PCR screening and replica plating on different selective allows to identify to correct clones.

**Figure 6 – figure supplement 2.**
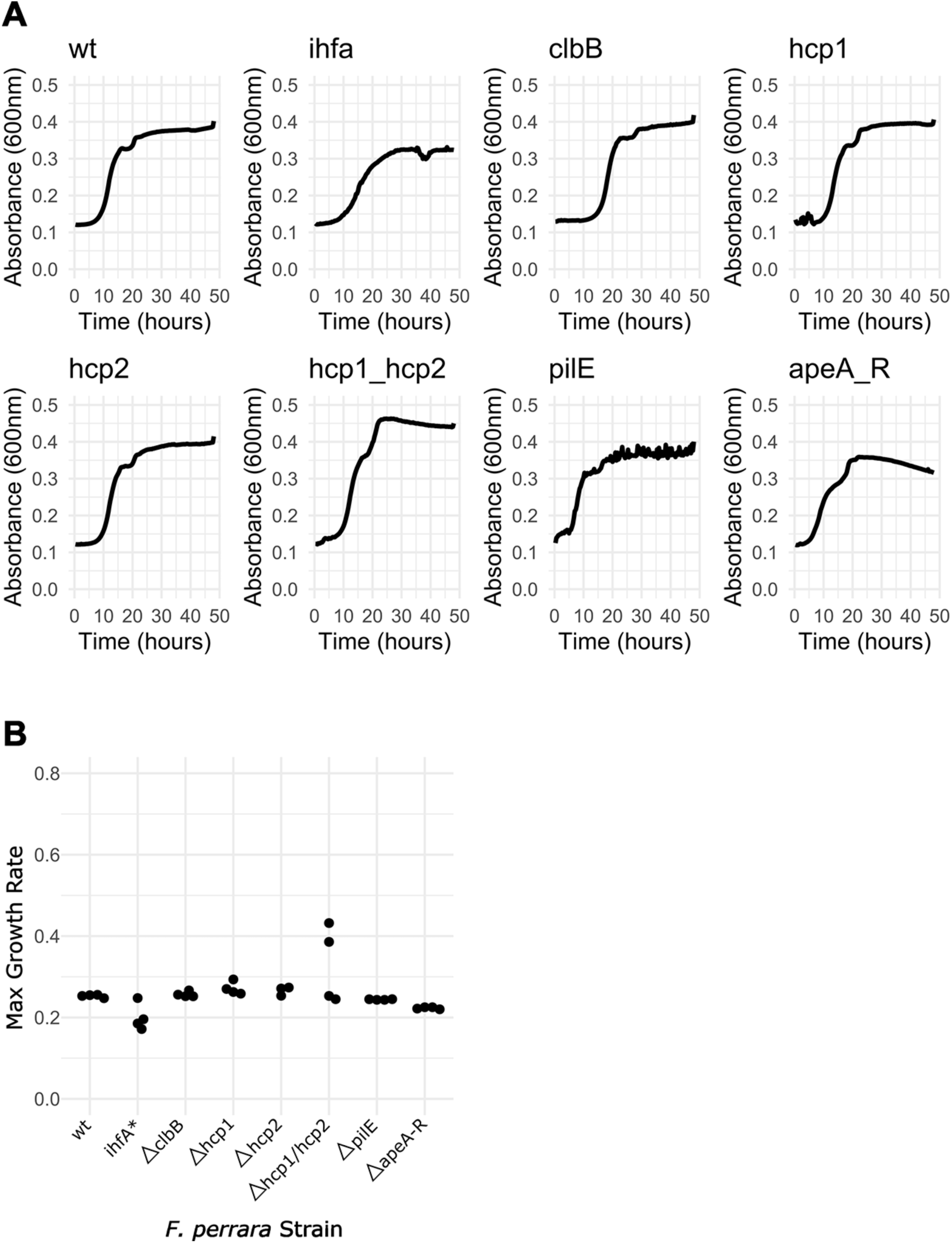
Growth Curves for the different *Frischella* strains. **(A)** The six gene deletion mutants, *ihfa\** and wt strains were diluted to OD_600_=0.05 and grown in BHI under anaerobic conditions at 35°C with continuous agitation. Absorbance was measured every 20 minutes for 48 hours. Per strain, four technical replicates were performed. **(B)** Max growth curve was calculated using the R package ‘growthcurver’. For the *hcp2* mutant, only three technical replicates were considered due to a contamination in one of the technical replicates.

**Figure 6 – figure supplement 3.**
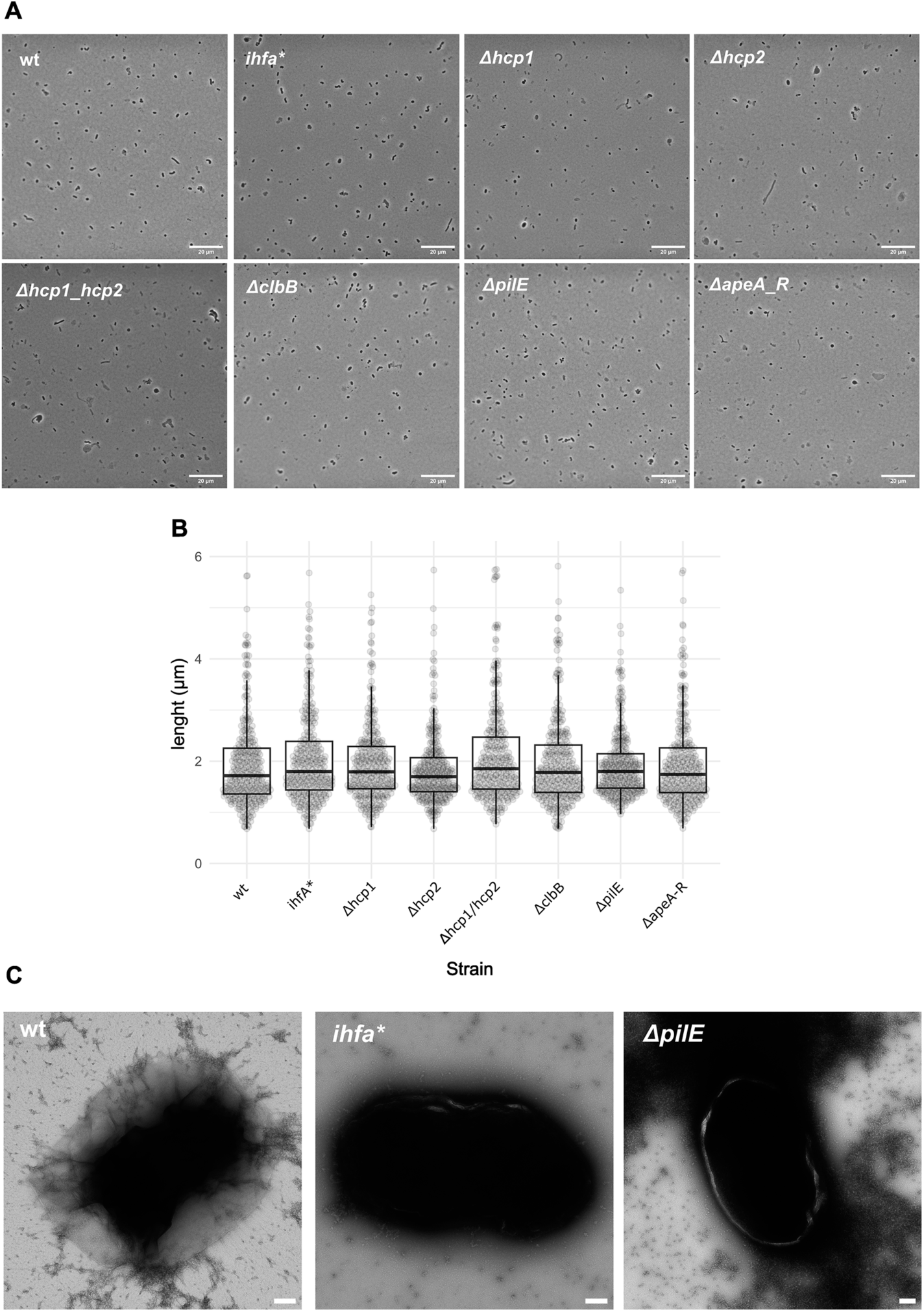
Single Cell imaging of *F. perrara* strains. **(A)** The six gene deletion mutants, *ihfa\**, and wt *F. perrara* strains were grown in liquid BHI, diluted to OD_600_= 0.1, plated in agar patches and imaged using a Nikon Ti inverted light microscope. Images were taken with a 100x objective. Scale bar indicates 20µm. **(B)** Cell length was quantified using the MicrobeJ plugin of ImageJ. **(C)** Electron Microscopy images were obtained for the wt, *ihfA\**, and *ΔpilE* strains. Scale bar indicates 200nm

**Figure 6 – figure supplement 4.**
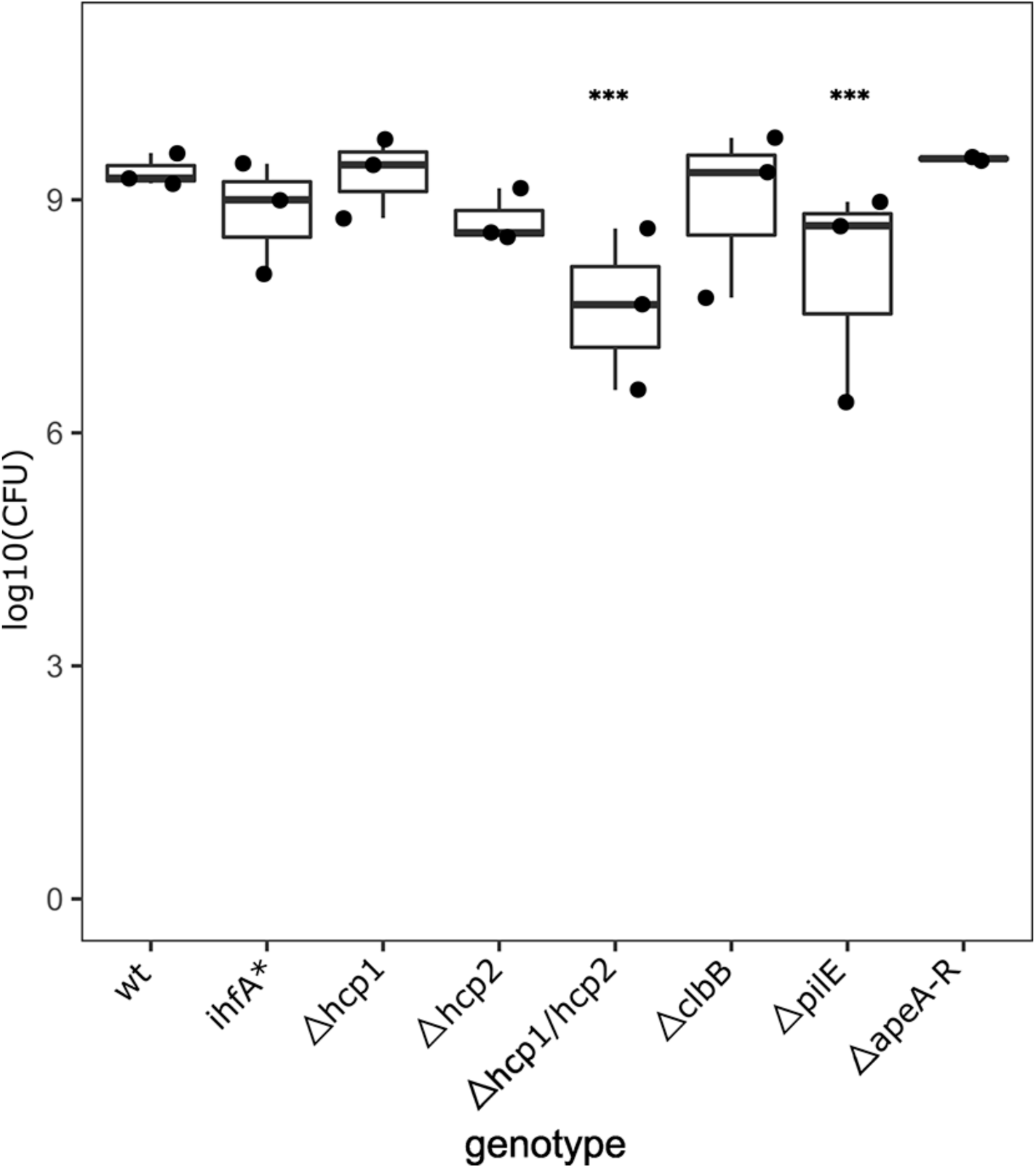
Correspondence between OD and CFU for *F. perrara* genotypes. For each *F. perrara* strain, a bacterial solution at OD_600_=0.1 was prepared, serially diluted and plated in BHIA medium. The number of CFUs present in 5µl of solution at OD_600_=0.1 was calculated based on the counts obtained from the serial dilutions. Statistics were calculated using a linear model with negative binomial distribution: *** p<0.0001, ** p<0.001.

**Figure 6 – figure supplement 5.**
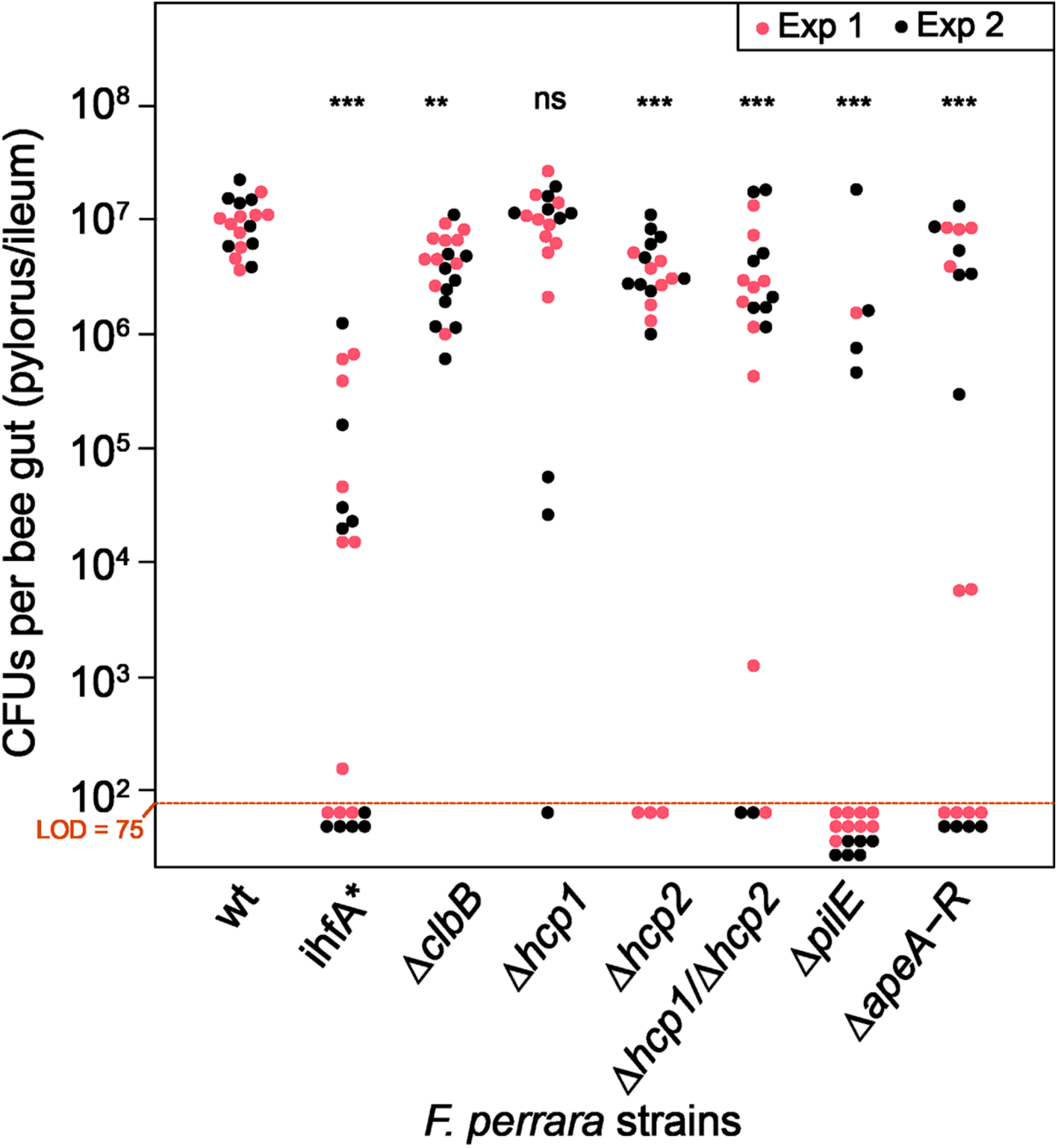
Same graph as shown in **Figure 6** but data points are labelled by experiment.

**Figure 6 – figure supplement 6.**
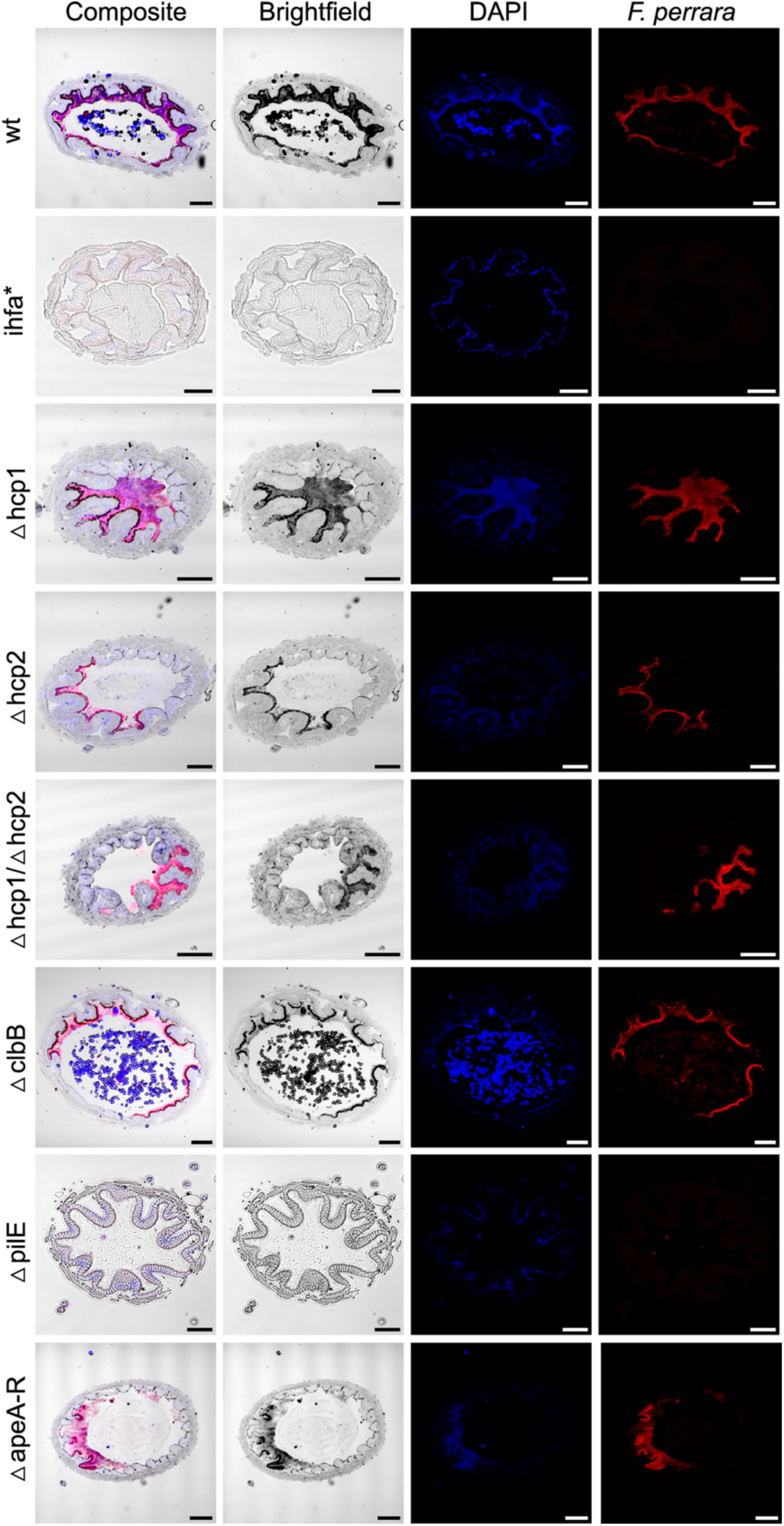
*F. perrara* colonization of the pylorus. Composite images are the same as in **Figure 6**. These were obtained by merging the brightfield, DAPI, and *F.perrara* probe individual images. Images were obtained with the 5x objective of the Zeiss LSM900. Hybridizations were done with probes specific for *F. perrara* (magenta). DAPI counterstaining of host nuclei and bacteria is shown in blue.

**Figure 6 – figure supplement 7.**
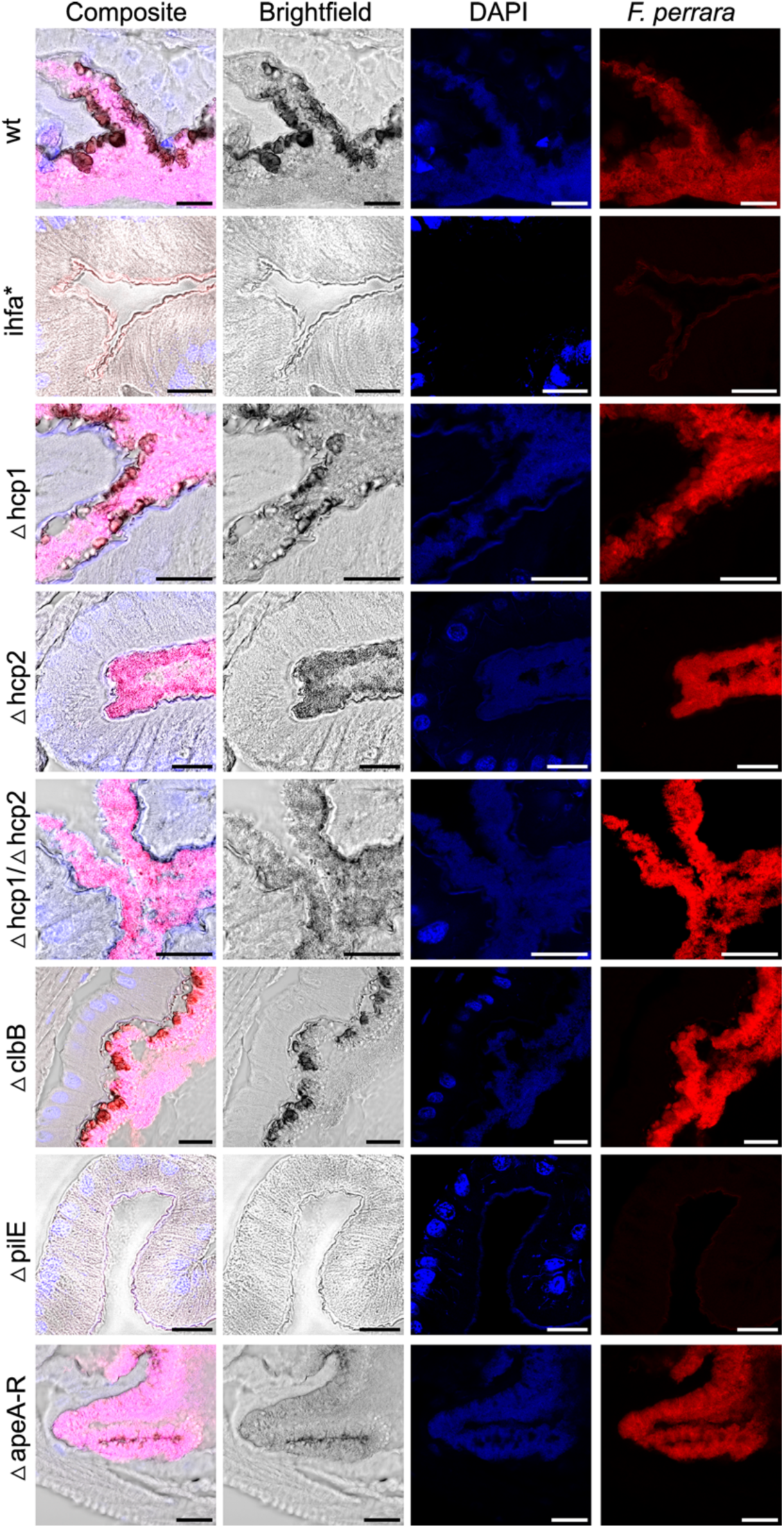
*F. perrara* colonization of the pylorus. Composite images are the same as in **Figure 6**. These were obtained by merging the brightfield, DAPI and *F.perrara* probe images. Images were obtained with the 40x objective of the Zeiss LSM900. Hybridizations were done with probes specific for *F. perrara* (magenta). DAPI counterstaining of host nuclei and bacteria is shown in blue.

**Figure 7 – figure supplement 1.**
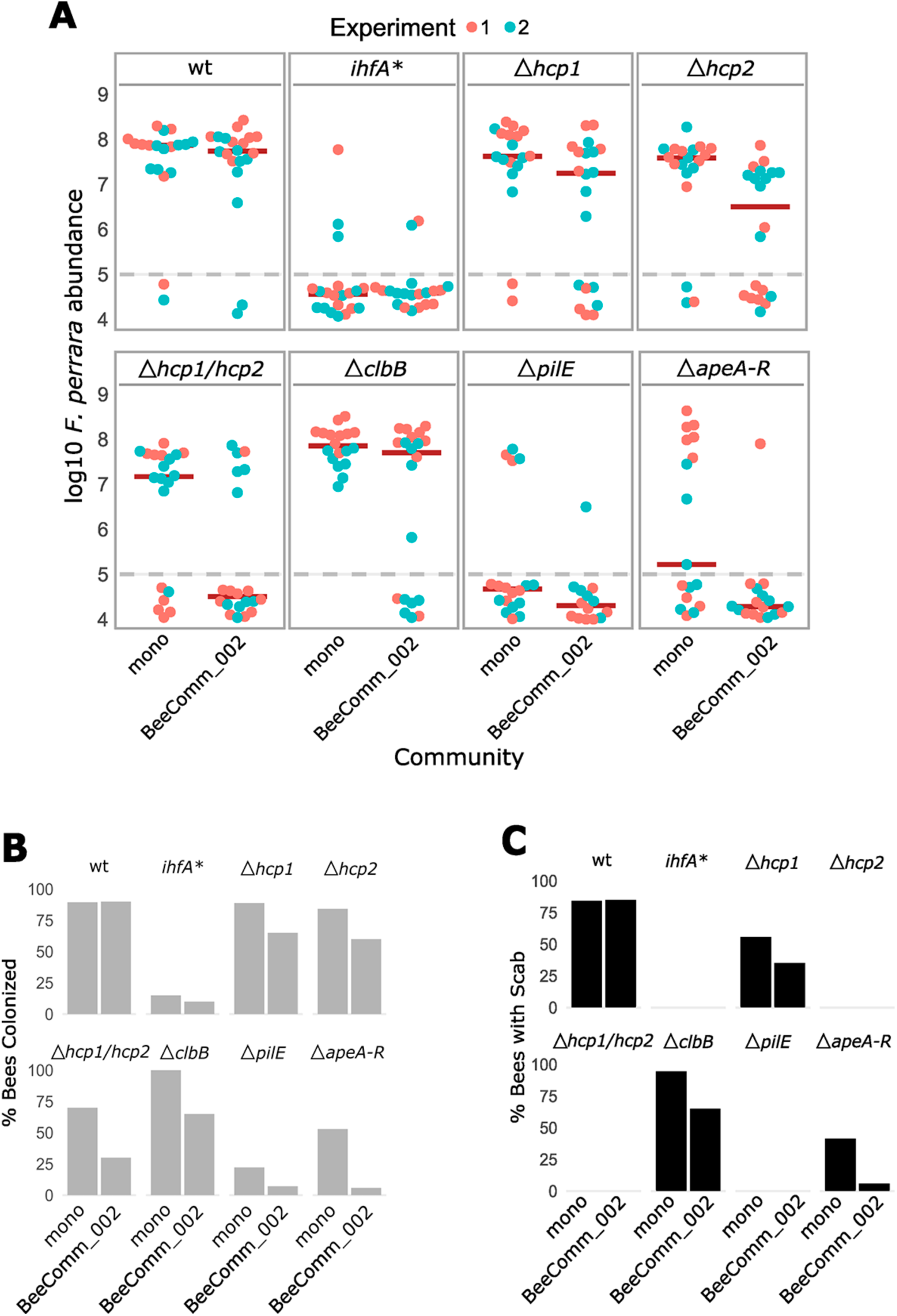
Colonization of wt, *ihfA*\* and gene-deletion mutants of *F. perrara*. **(A)** Same graph as shown in **Figure 7** but data points are labelled by experiment. **(B)** Percentage of bees that had colonization levels above the limit of detection. **(C)** Percentage of guts that had a scab.

## Supplementary Tables

**Supplementary Table 1.** Chemical shifts of enriched aryl polyene in DMSO–δ6 at 298 K. Key COSY and HMBC correlation of the enriched polyene.

| No | $\delta_C$ [ppm] | $\delta_H$ [ppm], mult. | coupling constant | integral |
| --- | --- | --- | --- | --- |
| 1 | 166.7 |  |  |  |
| 2 | 119.3 | 5.97, d | $J=15.1$ Hz | 0.5 |
| 3 | 144.6 | 7.33, dd | $J=11.6$ Hz<br>$J=15.1$ Hz | 1.2 |
| 4 | 129.5 | 6.47, dd overlap | $J=11.6$ Hz<br>$J=14.4$ Hz | 1.0 |
| 5 | 141.5 | 6.82, dd overlap | $J=11.4$ Hz<br>$J=14.5$ Hz | 0.9 |
| 6 |  |  |  |  |
| 7 |  |  |  |  |
| 8 |  |  |  |  |
| 9 |  |  |  |  |
| 10 |  |  |  |  |
| 11 |  |  |  |  |
| 12 |  |  |  |  |
| 13 |  |  |  |  |
| 14 |  |  |  |  |
| 15 | 128.7 | 7.22, d | $J=1.7$ Hz | 1.0 |
| 16 | 124.4 |  |  |  |
| 17 | 155.9 |  |  |  |
| 18 | 114.6 | 6.73, d | $J=8.3$ Hz | 1.2 |
| 19 | 125.2 | 7.13, dd | $J=1.9$ Hz<br>$J=8.3$ Hz | 1.1 |
| 20 | 15.6 | 2.11, s |  | 2.8 |
| 21 | 50.9 | 3.66, s overlap |  | 3.5 |

**Supplementary Table 2.** Strains used in this study.

| Species | Strain | Genotype* | Mutated gene | Reference | Parent strain |
| --- | --- | --- | --- | --- | --- |
| *F. perrara* | PEB0191/<br>ESL0157 | wild type (wt) | None | (31) | Primary isolate |
| *F. perrara* | ESL0158 | P83L (CCA→CTA) in *ihfA* | Fpe_00769 | This study | PEB0191 |
| *F. perrara* | ESL0910 | P82L (CCA→CTA) in *ihfB*;<br>Δ*apeO* | Fpe_00769;<br>Fpe_01928 | This study | PEB0191 |
| *F. perrara* | ESL0922 | L38S (TTA→TCA) in *ihfA*; $\Delta$ *pilE* * | Fpe_00769;<br>Fpe_00791*;<br>Fpe_01328 | This study | PEB0191 |
| *F. perrara* | ESL0854 | $\Delta$ *hcp1* * | Fpe_01742 | This study | PEB0191 |
| *F. perrara* | ESL0855 | Δ*hcp2* | Fpe_01947 | This study | PEB0191 |
| *F. perrara* | ESL0856 | Δ*hcp1*, Δ*hcp2* | Fpe_01742,<br>Fpe_01947 | This study | PEB0191 |
| *F. perrara* | ESL0888 | $\Delta$ *clbB* * | Fpe_00576 | This study | PEB0191 |
| *F. perrara* | ESL0921 | $\Delta$ *pilE* * | Fpe_01328 | This study | PEB0191 |
| *F. perrara* | ESL0957 | $\Delta$ *apeA-R* * | Fpe_01915-Fpe_01932 | This study | PEB0191 |

\*re-sequencing of the genomes revealed that some of the strains (indicated with an asterisk) carried an additional point mutation. However, these mutations were located in random genes and included synonymous as well as non-synonymous base substitutions unlikely to have an effect on the observed phenotypes. The genomic analysis of the re-sequenced strains can be found on the Switchdrive: https://drive.switch.ch/index.php/s/kCNTp4g7n60ffMi.

**Supplementary Table 3.** Primers used for qPCR quantification

| Primer name | Target | Sequence 5' to 3' | Tm (°C)<sup>a</sup> | Amplicon size (bp) | # Gene loci per genome | Standard curve |  |  | Reference |
| --- | --- | --- | --- | --- | --- | --- | --- | --- | --- |
|  |  |  |  |  |  | Primer Efficiency, R<sup>2</sup> | Slope, Intercept | LOD Cq<sup>b</sup> (#copies) |  |
| prPEN0013 | Actin AB023025 A.<br>*mellifera* | TGCCAACACTGT<br>CCTTTCTG | 58.4 | 156 | - | 1.896 = 89.6 %, 1.0 | -3.6, 37.57 | 33.7 (10) | (83) |
| prPEN0014 |  | AGAATTGACCCA<br>CCAATCCA | 56.4 |  |  |  |  |  |  |
| prLK-Frisch-042-F | *F. perrara* 16S<br>rRNA gene | GGAAGTTATGTG<br>TGGGATAAGC | 60.1 | 185 | 4 | 1.946 = 94.6 %, 0.995 | -3.45, 38.04 | 31.7 (100) | (18) |
| prLK-Frisch-043-R |  | CTATTCTCAGGTT<br>GAGCCCG | 60.5 |  |  |  |  |  |  |

a - Melting temperatures were calculated with the online tool described in (Kibbe 2007)
b - LOD refers to the limit of detection of primer sets, expressed here as the lowest number of plasmid copies reliably detected by qPCR when standard curves were performed

**Supplementary Table 4.**
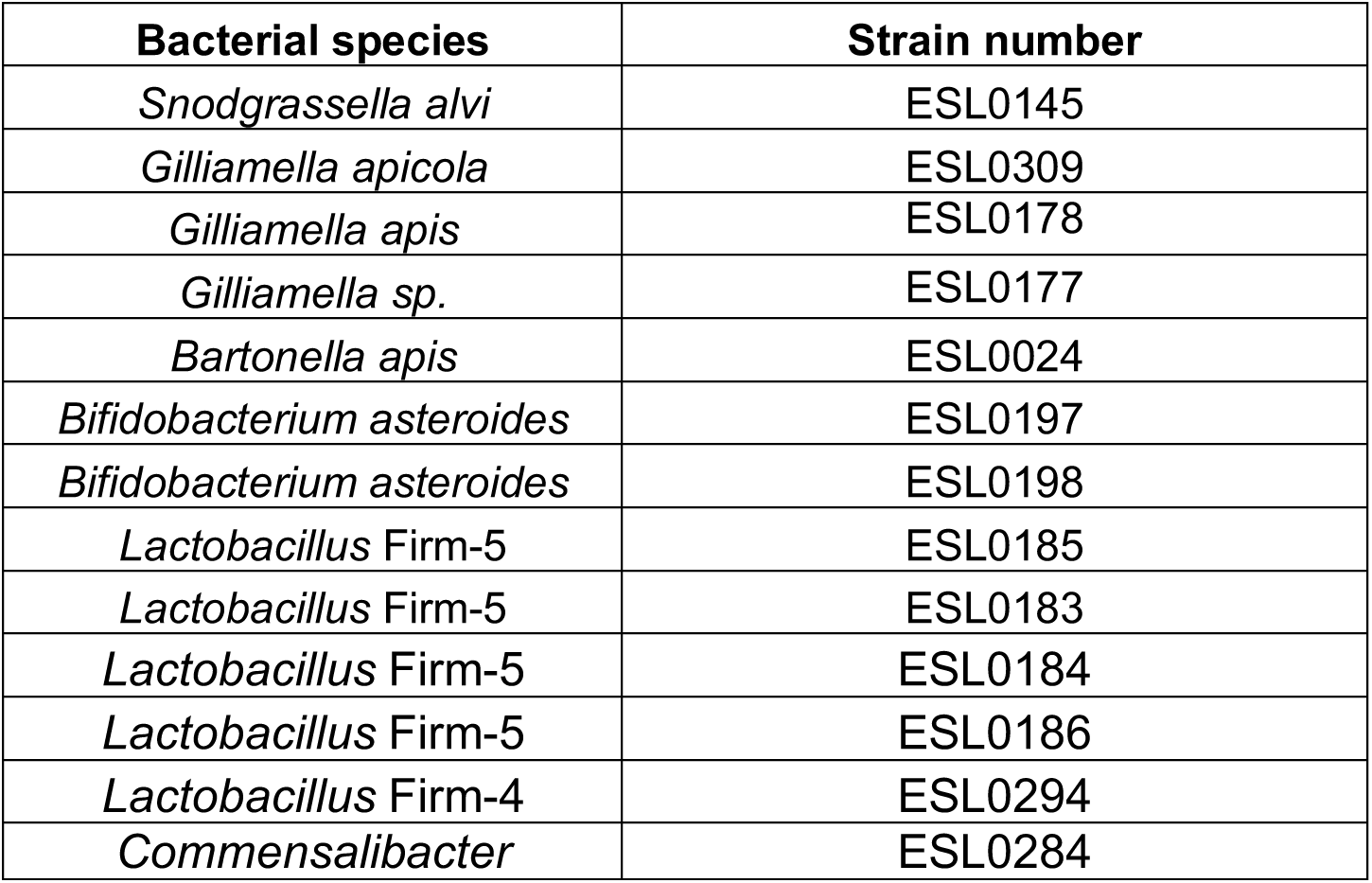
Bacterial strains included in the defined community.

